# Single-cell transcriptome analysis of CD34^+^ stem cell-derived myeloid cells identifies a CFU-GEMM-like population permissive to human cytomegalovirus infection

**DOI:** 10.1101/438457

**Authors:** Melissa Galinato, Kristen Shimoda, Alexis Aguiar, Fiona Hennig, Dario Boffelli, Michael A McVoy, Laura Hertel

## Abstract

Myeloid cells are important sites of lytic and latent infection by human cytomegalovirus (CMV). We previously showed that only a small subset of myeloid cells differentiated from CD34^+^ hematopoietic stem cells is permissive to CMV replication, underscoring the heterogeneous nature of these populations. The exact identity of susceptible and resistant cell types, and the cellular features characterizing permissive cells, however, could not be dissected using averaging transcriptional analysis tools such as microarrays and, hence, remained enigmatic. Here, we profile the transcriptomes of ∼ 7000 individual cells at day one post-infection using the 10X genomics platform. We show that viral transcripts are detectable in the majority of the cells, suggesting that virion entry is unlikely to be the main target of cellular restriction mechanisms. We further show that viral replication occurs in a small but specific sub-group of cells transcriptionally related to, and likely derived from, a cluster of cells expressing markers of Colony Forming Unit – Granulocyte, Erythrocyte, Monocyte, Megakaryocyte (CFU-GEMM) oligopotent progenitors. Compared to the remainder of the population, CFU-GEMM cells are enriched in transcripts with functions in mitochondrial energy production, cell proliferation, RNA processing and protein synthesis, and express similar or higher levels of interferon-related genes. While expression levels of the former are maintained in infected cells, the latter are strongly down-regulated. We thus propose that the preferential infection of CFU-GEMM cells may be due to the presence of a pre-established pro-viral environment, requiring minimal optimization efforts from viral effectors, rather than to the absence of specific restriction factors. Together, these findings identify a potentially new population of myeloid cells susceptible to CMV replication, and provide a possible rationale for their preferential infection.

**AUTHOR SUMMARY:** Myeloid cells such as monocytes and dendritic cells are critical targets of CMV infection. To identify the cellular factors that confer susceptibility or resistance to infection, we profiled the transcriptomes of ∼ 7,000 single cells from a population of semi-permissive myeloid cells infected with CMV. We found that viral RNAs are detectable in the majority of the cells, but that marked expression of CMV lytic genes occurs in only a small subset of cells transcriptionally related to a cluster of CFU-GEMM progenitors that express similar amounts of transcripts encoding interferon-related anti-viral factors as the rest of the population but higher levels of transcripts encoding proteins required for energy, RNA, and protein production. We thus conclude that the preferential infection of CFU-GEMM cells might be due to the pre-existing presence of an intracellular environment conducive to infection onset, rather than to the absence of anti-viral factors restricting viral entry or initial gene expression. Together, these findings uncover a new type of myeloid cells potentially permissive to CMV infection, expand our understanding of the cellular requirements for successful initiation of CMV infection, and provide new pro- and anti-viral gene candidates for future analyses and therapeutic interventions.

## INTRODUCTION

Infection by human cytomegalovirus (CMV) is common and usually asymptomatic in healthy individuals, but can be the source of serious disease in hosts with naïve or compromised immune functions such as fetuses, newborns, AIDS patients, and solid organ or bone marrow transplant recipients [1, 2]. CD34^+^ hematopoietic stem cells (HSC) and derived monocytes, macrophages and dendritic cells are important sites of CMV latency and reactivation, as well as of lytic infection *in vivo* (for recent reviews, see [3–6]). CMV interactions with these cells have thus been intensively studied, using a variety of different cell culture models [7–11].

We previously showed that exposure of cord blood CD34^+^ HSC to specific cytokines such as Flt3 ligand (FL) and transforming growth factor β1, known to instruct their differentiation into Langerhans cells [12, 13], gives rise to myeloid cell populations that are semi-permissive to CMV infection. While only 2-3% of non-activated cells obtained at the end of the differentiation period allowed expression of the viral immediate early 1 and 2 (IE1/IE2) proteins, which are essential for infection onset and progression, cell activation by exposure to granulocyte-macrophage colony-stimulating factor (GM-CSF), fetal bovine serum (FBS), CD40 ligand (CD40L) and lipopolysaccharide (LPS) partially released this initial block, raising the proportion of IE1/IE2^+^ cells by 5-10 fold [14–16]. Unexpectedly, however, non-activated cells produced higher yields per IE1/IE2^+^ cell than activated cells, suggesting that signaling by GM-CSF, FBS, CD40L and LPS may trigger the establishment of a second block to infection progress, acting after IE gene expression and negatively impacting viral progeny production. This second block is unlikely to be due to defects in progeny release, as non-activated and activated cells generated similar ratios of cell-free to cell-associated virus [16], but may instead depend on impairments in the assembly of viral replication compartments (Galinato and Hertel, unpublished).

Because of their ability to restrict infection progress at multiple steps of the viral replication cycle, activated myeloid cells represent an outstanding model to study the determinants of cellular susceptibility to CMV infection. Their intrinsic heterogeneity, however, thus far precluded the identification of cellular factors supporting or restricting infection using averaging gene expression analysis tools such as microarrays. Here, we took advantage of the most recent developments in single-cell RNA sequencing technologies to provide the first transcriptional profiling of CMV-infected, activated myeloid cells conducted at the single-cell level, and the first comparison of gene expression changes occurring in infected and bystander cells co-existing in the same population.

We show that: 1) more than half of the cells contain detectable viral transcripts at day one post-infection, with only a small minority (∼ 2%) displaying an expression pattern consistent with progression to lytic replication. This indicates that restrictions to viral entry may contribute to, but are not the main determinant of resistance; 2) lytically-infected cells are transcriptionally related to a specific cluster of bystander cells with the hallmarks of CFU-GEMM, suggesting that this type of cells may be a previously unidentified target of CMV lytic infection; 3) compared to the remainder of the population, CFU-GEMM cells express similar or higher levels of IFN-related genes with anti-viral roles, which are strongly down-regulated in infected cells, indicating that CFU-GEMM cells are not defective in their ability to recognize and respond to CMV infection; 4) also compared to the remainder of the population, CFU-GEMM cells are enriched in transcripts encoding proteins involved in mitochondrial energy production, S-phase control, and RNA and protein production. Expression levels of these genes remain largely unchanged in lytically-infected cells, suggesting that that preferential infection of CFU-GEMM cells is likely due to the presence of a transcriptional landscape already optimized for viral replication, and requiring little conditioning effort from viral effectors, rather than to an intrinsic inability to recognize and respond to the presence of viral products.

Together, these data identify a new myeloid cell type potentially permissive to CMV replication, broaden our knowledge of the cellular determinants of susceptibility to infection, and reveal the identity of new pro- and anti-viral factors involved in regulating CMV tropism for myeloid cells.

## RESULTS

### CD34^+^ HSC-derived myeloid cell populations are semi-permissive to CMV infection

To identify cellular factors potentially involved in regulating the susceptibility of myeloid cells to CMV infection, we sought to analyze the transcriptome of permissive and non-permissive cell types derived from the differentiation of CD34^+^ HSC *in vitro*. To select a representative population, the CD34^+^ HSC isolated from the cord blood of twelve different donors were separately cultured in the presence of a cytokine cocktail known to promote the development of Langerhans-type dendritic cells [12, 13]. Differentiated cells were then activated by exposure to GM-CSF, FBS, CD40L and LPS, and infected with CMV strain TB40/E. Consistent with our previously published data [14–16], cell numbers did not increase over time (not shown), and only 3 ± 1.5 % of non-activated but 10 ± 5 % of activated cells expressed the viral IE1/IE2 proteins at day two post-infection (pi), notwithstanding the use of a multiplicity of infection (MOI) of ten pfu/cell, which is sufficient to infect the totality of permissive cell types such as fibroblasts (Fig 1A and B). Despite containing higher numbers of IE1/IE2^+^ cells at each time point, activated cell populations produced lower intracellular progeny amounts per IE1/IE2^+^ cell than non-activated cells (Fig 1C and D).

**Figure 1.**
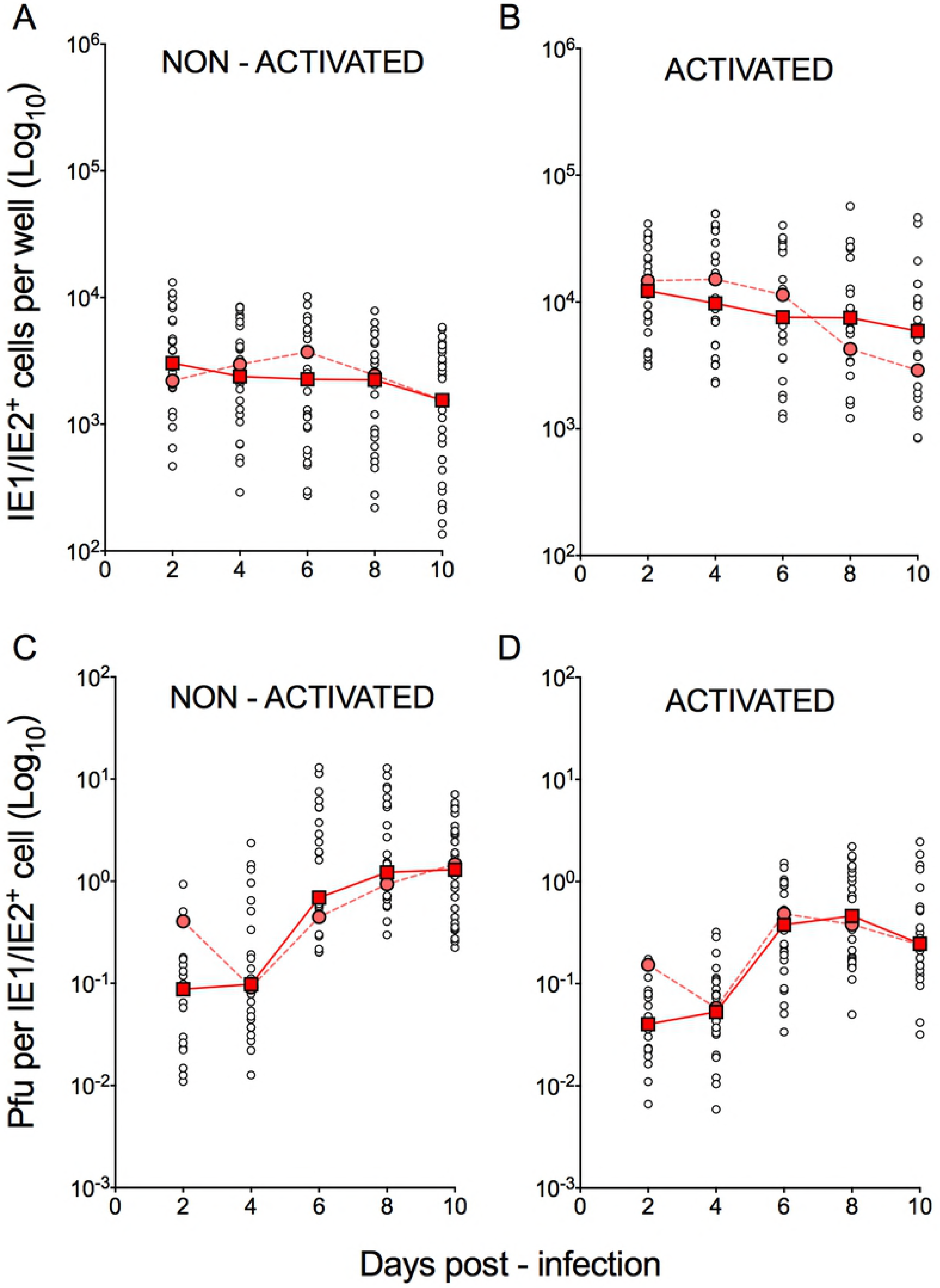
Susceptibility to CMV infection of non-activated and activated myeloid cell populations differentiated from cord blood CD34^+^ HSC from different donors. Non-activated and activated myeloid cell populations differentiated from cord blood CD34^+^ HSC from twelve different donors were exposed to CMV strain TB40/E at an MOI of ten and analyzed at the indicated days pi. A small portion of each population was subjected to immunofluorescence staining to determine the number of IE1/IE2^+^ cells present in each well (A-B), while the remainder of the cells were sonicated and used in titration assays to quantify intracellular progeny yields (C-D). Open circles represent data from individual donors; red squares indicate median values at each time point; salmon circles depict data obtained from the CD34^+^ HSC of representative donor 113G.

Activated cells differentiated from the CD34^+^ HSC of donor 113G (Fig 1, pink circles) were then selected as representative, and subjected to single-cell RNA sequencing at day one using the 10X Genomics Chromium platform [17]. Activated cells were chosen to ensure data collection from sufficient numbers of infected cells, and to facilitate the identification of potential cellular inhibitors of viral replication, whereas the day one time point was selected to allow sufficient time for infection to start, while limiting the extent of virus-induced changes to the cellular transcriptional landscape. A median of 2,305 genes and 10,627 transcripts were detected in the 6,837 cells profiled, and the total number of genes with at least one count in any cell was 20,899. After reduction by principal components analysis, data was visualized in two dimensions using the *t*-distributed stochastic neighbor embedding (*t*-SNE) algorithm [18], which displays cells with similar transcriptional profiles as nearby points, and cells with dissimilar transcriptional profiles as distant points with high probability (Fig 2A). Cells thus represented on *t*-SNE plots were then interrogated for specific gene transcripts using Loupe™ Cell Browser [19].

**Figure 2.**
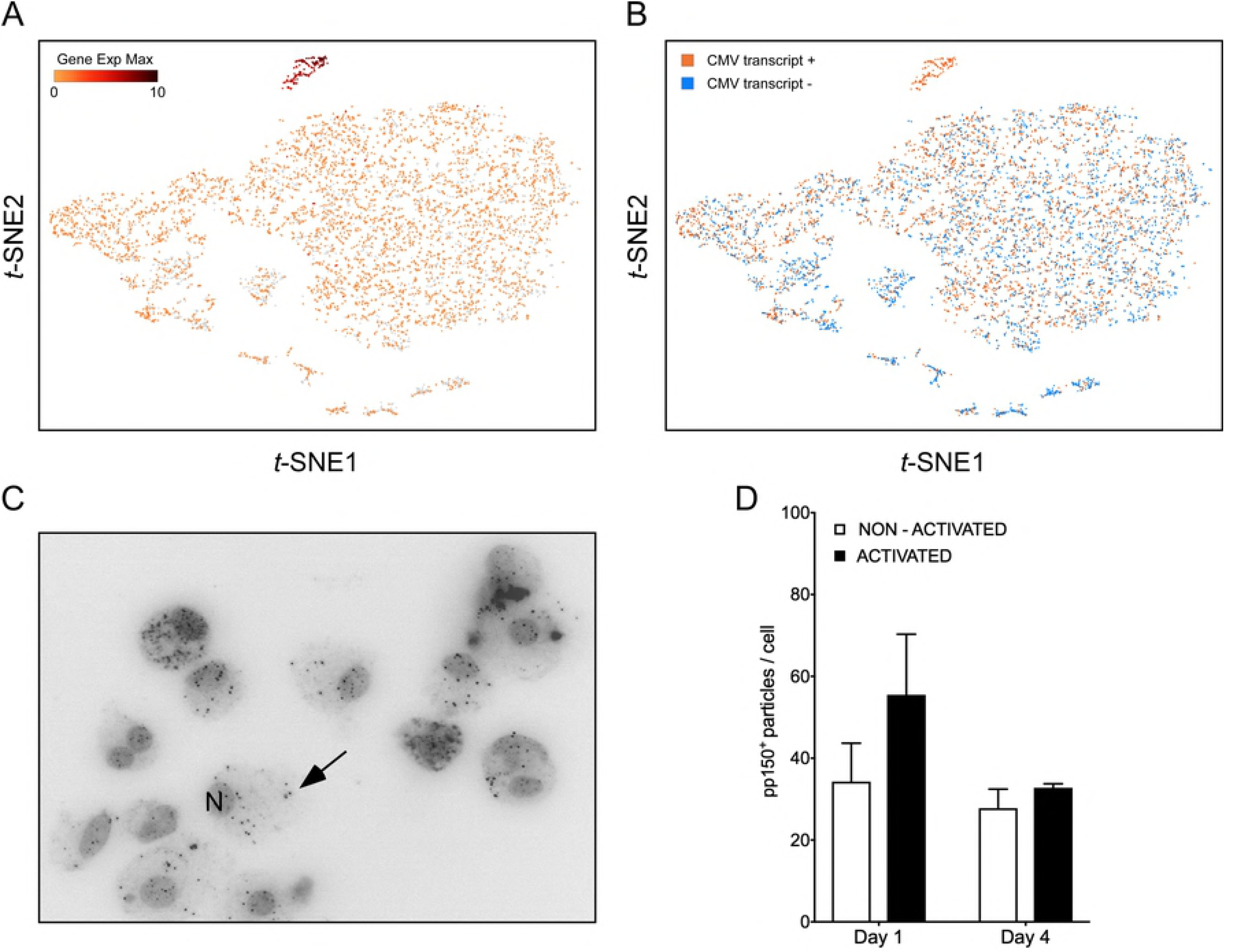
Detection of viral transcripts in a large proportion of cells in the population. (A-B) *t*-SNE projections of data from each of the 6,837 profiled cells depicted as dots colored based on their content in transcripts mapping to viral ORFs and displayed in a quantitative (A) or qualitative (B) manner. In A, cells lacking detectable viral transcripts are shown in grey, while cells containing viral RNAs are colored in shades of red depending on transcript amounts (Log_2_ Gene Exp Max). In B, cells containing or lacking viral transcripts are shown in orange and blue, respectively. (C) Merged representative image of activated myeloid cells infected with TB40/E at an MOI of ten, harvested at day one pi, and stained for pp150, to visualize viral particles (arrow), and with Hoechst 33342 to visualize cell nuclei (N). (D) Number of pp150^+^ particles per cell at day one and day four pi, manually counted from 5-15 images/sample of non-activated and activated myeloid cells in four independent experiments. Median and median absolute deviation values are shown.

### Viral transcripts are detected in the majority of the cells, but their presence is not associated with expression of specific cellular genes

Query of the *t*-SNE projection data for the presence of viral RNA revealed that 59% of the cells in the population contained at least one viral transcript (Fig 2A and B). RNAs mapping to the viral open reading frames (ORFs) UL4/UL5, US34, UL145 and UL16/17, and to the non-coding RNA2.7 and RNA1.2, were present in the largest number of cells, accounting for half of the CMV-transcript^+^ population. These viral RNA^+^ cells were dispersed throughout the entire population, suggesting that infection had occurred into the majority of the cells. However, to avoid introducing perturbations potentially affecting cellular transcription, non-penetrated viral particles were not enzymatically removed from the cell surface. Consequently, some of the detected transcripts may have originated from virions still attached to the outside of cells or from penetrated capsids that did not reach the nucleus. The specific viral RNAs that were detected in the largest proportion of the cells, however, are not amongst those reported to be packaged into virions [20–22]. Moreover, more than half of the 26 transcripts found in > 200 cells mapped to viral ORFs known to be expressed with immediate-early or early kinetics (not shown), suggesting that they were likely newly synthesized from the viral genome.

Staining of infected cells for the capsid-associated phosphoprotein pp150 [14, 23, 24] (Fig 2C) also revealed the presence of viral particles associated with 55 ± 15 % of activated cells and in 34 ± 9 % of non-activated cells at day one pi (Fig 2D). These results are consistent with our previously published data using CMV strain TB40-BAC4, a BAC-cloned variant of TB40/E [14], although, in contrast to TB40-BAC4, TB40/E virions remained visible on, or within, the cells until at least day four pi.

To identify cellular factors potentially involved in restricting viral entry, the gene expression profile of CMV-transcript^+^ cells was compared to that of CMV-transcript^−^ cells. Only five cellular genes scored as differentially expressed between the two groups of cells with P < 0.05, but none was present in the totality, nor in the majority, of CMV-transcript^−^ or CMV-transcript^+^ cells (S1 Dataset). The two genes expressed in the largest number of cells, RETN and TJP1 (for gene names, see S2 Table 1), were detected in only 303 and 124 cells, respectively, and were distributed in both populations: RETN was found in 153 CMV-transcript^+^ vs. 150 CMV-transcript-cells, and TJP1 in 104 CMV-transcript^+^ vs 20 CMV-transcript-cells.

The extent of expression of genes encoding potential CMV entry receptors, such as EGFR [25–28], PDGFRA [29–31], THY1/CD90 [32, 33], the integrins αVβ3, α2β1, and α6β1 [34–36], and BSG [37] was also queried. EGFR, THY1/CD90, and integrins β3, α2, and α6 were either not expressed at all or were found in less than ten cells, while PDGFRA was expressed in only 356 cells, and then only at low levels. Integrin β1 and BSG, by contrast, were present in larger numbers of cells (2568 and 5810, respectively), but these did not preferentially segregate with the CMV-transcript^+^ group.

Together, these findings indicate that viral entry is unlikely to be the main roadblock restricting infection onset, that cells devoid of viral transcripts do not transcribe specific factor(s) restricting virion entry, and that cells containing viral RNAs do not selectively express genes encoding entry facilitators, including surface molecules reported to act as CMV entry receptors in other cell types.

### Transcription of viral lytic genes proceeds in a small group of cells lacking expression of select cellular genes

Eleven genetic loci were identified as required for efficient CMV genome replication in transient co-transfection replication assays [38, 39]. These encode the transcriptional activators/regulators IE1, IE2, UL112/113, UL84 and IRS1/TRS1, the anti-apoptotic factors UL36-38, and six members of the viral DNA replication complex, i.e. the DNA polymerase UL54, the polymerase accessory factor UL44, the helicase UL105, the primase UL70, the primase associated factor UL102, and the single-stranded DNA binding protein UL57. To identify cells ostensibly progressing towards lytic replication, the population was queried for the presence of viral transcripts encoding each of these proteins.

A total of 278 cells, corresponding to ∼ 4% of the entire population and ∼ 7% of CMV-transcript^+^ cells were UL122^+^ (IE2) and/or UL123^+^ (IE1), and 42% of these expressed both. These proportions were in agreement with those obtained by immunofluorescence staining of infected cells from donor 113G harvested at day one pi (3.6% IE1/IE2^+^, data not shown).

Consistent with progression towards lytic replication (Fig 1D), UL122^+^/UL123^+^ cells were also found to express transcripts encoding UL112/113 (91% triple-positive), UL84 (77%), IRS1/TRS1 (59%), UL36 (84%), UL37 (13%), UL38 (83%), and three replication complex components, namely UL54 (64%), UL105 (70%), and UL102 (56%) (Fig 3A). By contrast, RNAs corresponding to UL44, UL70, and UL57 were not detected. Staining of infected cells confirmed that the UL84 protein was present in 37 ± 19% of activated, IE1/IE2^+^ cells at day two pi (Fig 3B and E). Interestingly, while the UL44 and UL57 proteins were also observed at day two pi (Fig 3C and D), the proportion of IE1/IE2^+^ cells co-expressing each of these polypeptides in activated cells was significantly lower than in non-activated cells (Fig 3E). This suggests that assembly of the viral DNA replication complex in activated cells might be impaired, perhaps on account of delayed or inefficient transcription of specific complex members.

**Figure 3.**
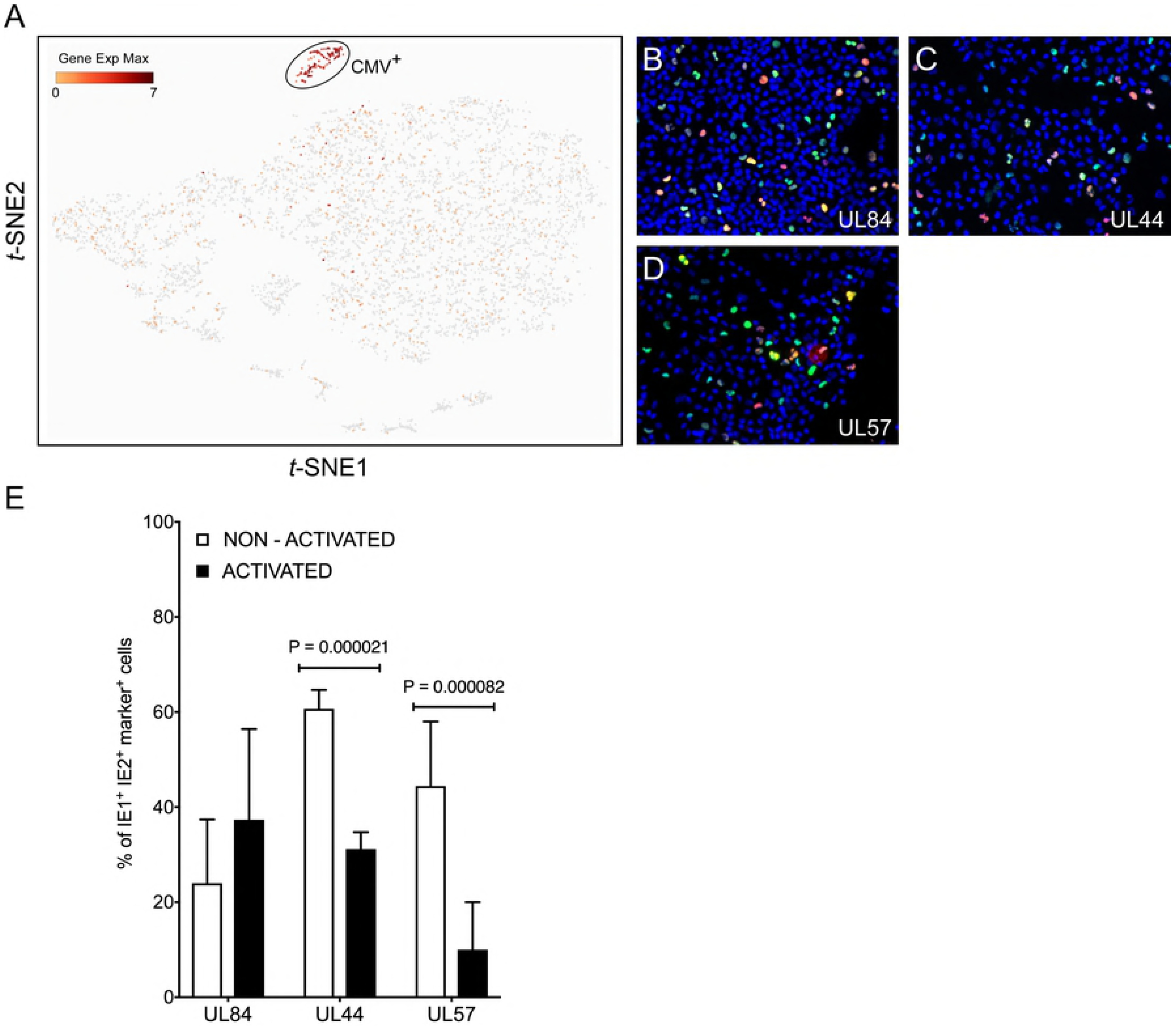
Detection of viral transcripts associated with progression toward lytic replication in a specific subset of cells. (A) *t*-SNE projection of data from profiled cells colored based on their quantitative (Log_2_ Gene Exp Max) cumulative content in transcripts mapping to the viral RNAs UL122, UL123, UL112/113, UL84, TRS1, UL36, UL37, UL38, UL54, UL44, UL102, UL105, UL70 or UL57. (B-D) Merged representative images of activated myeloid cell populations infected with TB40/E at an MOI of ten, harvested at day two pi, and co-stained for IE1/IE2 (green) plus UL84 (red, B), or UL44 (red, C) or UL57 (red, D) and with Hoechst 33342 (blue) to visualize cell nuclei. Cells expressing both IE1/IE2^+^ and the marker protein of interest appear yellow. (E) Percentage of IE1/IE2^+^ cells co-expressing the UL84, UL44, or UL57 proteins at day two pi, manually counted from 5-15 images/sample of non-activated and activated myeloid cells in seven independent experiments. Median, median absolute deviation and P values from unpaired T tests are shown.

The data from cells containing the above viral transcripts, plus several others, comprised a tight and well separated cluster of 138 points on the *t*-SNE projection, which we collectively named CMV^+^ (Fig 3A). Comparison of the transcriptional profile of CMV^+^ and CMV^−^ cells identified 629 genes as being more highly expressed in CMV^−^ cells, but none was associated with significant P values (< 0.05), nor was present in the totality of CMV^−^ and absent in CMV^+^ cells. Sixty cellular genes had more than four-fold higher mean expression levels in CMV^−^ than in CMV^+^ cells, with nine being present in more than 50% of CMV^−^ cells but less than 50% of CMV^+^ cells (S3 Dataset and S4 Fig). Encouragingly, and as expected for virus-exposed cells, five of these nine genes encoded well-known type I interferon (IFN)-inducible proteins involved in mediating innate immune responses to viruses, i.e. MX1, OAS1, OAS2, IFIT3, and USP18. In line with activated myeloid cells, the majority of CMV^−^ cells also expressed the TNF-α and LPS-inducible protein CYTIP [40], and the CD40 ligand-inducible costimulatory molecule CD80 [41], plus two genes, one coding for the orphan G protein-coupled receptor GPR157 and one for DUSP4, whose transcriptional regulation by viruses or other stimuli has not been assessed.

Together, these data indicate that although CMV transcripts are associated with the majority of cells, lytic infection proceeds in only a small sub-population containing lower amounts of a handful of genes, most of which encode known powerful antiviral proteins.

### CMV^+^ cells are closely related to a specific sub-cluster of cells within the population

The direct comparison of CMV^+^ and CMV^−^ cells failed to yield transcripts with statistically significant differences in expression levels between the two groups, suggesting that resistance to infection is unlikely to be conferred by a single set of “universal” restriction factors highly expressed in all CMV^−^ cells but absent in CMV^+^ cells. Rather, we hypothesized that the CMV^−^ population might be comprised of several different cell types, each resisting infection due to the expression of anti-viral genes, or to the lack of pro-viral factors, specific to each sub-group.

Cell clustering using the K-means algorithm indeed revealed the presence of multiple different sub-groups of cells within the population, each characterized by distinct transcriptional profiles (Fig 4A). To identify the cluster most closely related to CMV^+^ cells, the mean number of transcripts/cell for each gene in the CMV^+^ cluster was divided by the mean number of transcripts/cell for each gene in each of the other nine clusters, and the frequency distribution of all Log_2_ ratios was plotted. A nonlinear regression fit test using the least squares method revealed that all distributions could be described by the Gaussian function, and that the histogram with the mean value closest to zero (0.184), the smallest standard deviation (0.814), and the highest R^2^ value (0.995) belonged to the CMV^+^ versus cluster 6 comparison (Fig 4B). A Wilcoxon signed rank test also identified the Log_2_ CMV^+^/cluster 6 ratio distribution as the one whose median values differed the least from zero, suggesting that the transcriptional profiles of CMV^+^ and cluster 6 cells were the most similar to each other.

**Figure 4:**
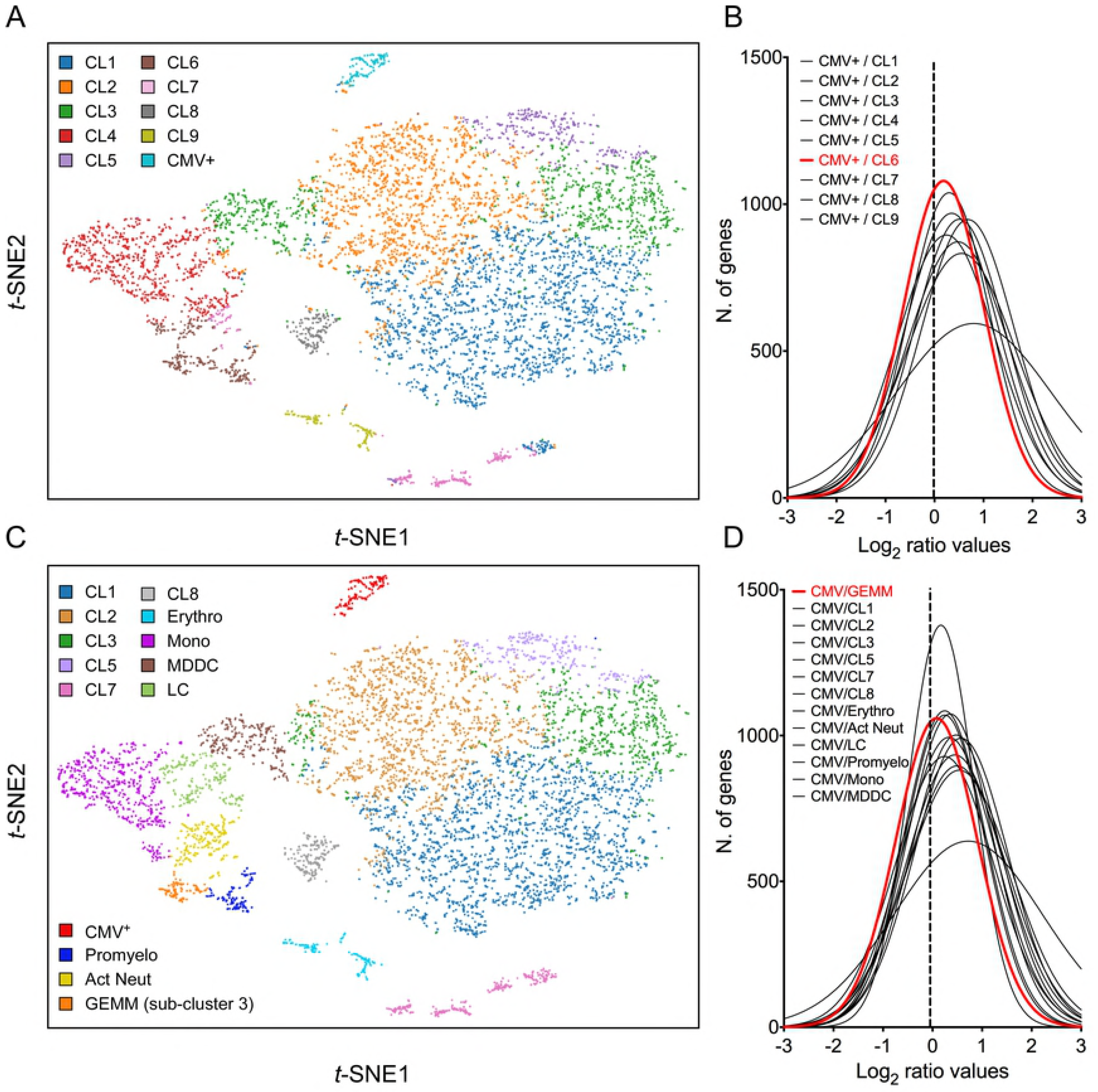
Identification of cluster 6 as the most closely related to the CMV^+^ cluster. (A) *t*-SNE projection of data from profiled cells partitioned into clusters by the K-means clustering algorithm using *K* = 10. (B) Distribution of Log_2_ ratio values obtained by dividing the mean number of transcripts/cell for each gene in the CMV^+^ cluster by the mean number of transcripts/cell for each gene in each of the other nine clusters. The distribution with the Log_2_ mean value closest to zero (dotted vertical line) is depicted by a red line. (C) *t*-SNE projection of data from profiled cells partitioned into clusters by the K-means clustering algorithm using *K* = 10, and further sub-divided based on marker gene expression. (D) Distribution of Log_2_ ratio values, as described in (B), comparing the CMV^+^ cluster the other 13 clusters. CL = cluster; Erythro = erythrocytes-megakaryocytes; Mono = monocytes; MDDC = monocyte-derived dendritic cells; LC = Langerhans cells; Promyelo = promyelocytes; Act Neut = activated neutrophils; GEMM = colony-forming unit-granulocyte, erythrocyte, monocyte/macrophage, megakaryocyte.

To uncover the identity of cluster 6 cells, the genes most selectively expressed by this cluster relative to the rest of the population were identified using the 10X Genomics Cell Ranger software [42], and their expression range *in vivo* was assessed using publicly available gene expression databases and literature data. Sixty-nine genes were selected as being highly differentially expressed (Log_2_ cluster 6/rest of the cells fold change > 4, P values < 10^−15^, S5 Dataset). Seven of these (ELANE, PRTN3, AZU1, MPO, PRSS57, CTSG and RNASE2), coding for markers of neutrophil precursors [43, 44] were predominantly or exclusively expressed in a sub-group of 116 cells, which we designated “promyelocytes” (S6 Fig A). Four more genes (RETN, S100A8, S100A9 and S100A12), encoding proteins secreted by activated neutrophils under pro-inflammatory conditions [45, 46], were abundant in promyelocytes and in a separate group of ∼ 205 cells, designated “activated neutrophils” (S6 Fig B). The remaining 57 genes encoded mostly DNA replication and cell cycle regulators, and were present, either exclusively or overlapping with promyelocytes and CMV^+^ cells, in a third sub-group of 93 cells (S6 Fig C), designated “sub-cluster 3”.

In addition to separating cluster 6 into three sub-clusters, other groups of cells were identified based on their expression of known markers such as CD14 and CD68 (monocytes), CD207/langerin and CD1a (Langerhans cells), CD1b (monocyte-derived dendritic cells), and hemoglobins (erythrocytes) (Fig 4C). Of note, and in keeping with CD34^+^ HSC differentiation towards myeloid (rather than lymphoid) lineages, no T or B cell specific transcripts were found.

While none of the promyelocyte- and activated neutrophil-specific genes were also expressed by CMV^+^ cells, 18 (32%) of sub-cluster 3 marker genes were shared, some almost exclusively, with the CMV^+^ group. This suggested that sub-cluster 3 cells in specific might be related to the CMV^+^ cell cluster. To further verify this, the mean number of transcripts/cell for each gene in the CMV^+^ cluster was divided by the mean number of transcripts/cell for each gene in each of the other 13 clusters, and the frequency distribution of Log_2_ ratios was plotted. The histogram whose median value differed the least from zero did indeed correspond to the CMV^+^ versus sub-cluster 3 comparison (Fig 4D), confirming that the transcriptional profile of these two groups are the most closely related.

### Sub-cluster 3 is comprised of cells with CFU-GEMM hallmarks

To more precisely identify the cell type comprising sub-cluster 3, the list of 115 genes more abundantly (average transcript count > 0.3) and most differentially (Log_2_ fold change > 3, P < 0.0005) transcribed in these cells relative to all other clusters was compared to gene lists from two recently published single-cell analyses of human hematopoiesis [47, 48] (S7 Dataset). Seventy-two transcripts were found among the list of genes reported to be differentially expressed in 16 discrete bone marrow populations by Velten L. et al. [48], with the largest proportion falling within the “G2/M phase” (56%) and the “Immature myeloid progenitors with high cell cycle activity” (24%) categories. A total of 108 genes were also found among the transcripts classified as differentially expressed in seven human cord blood populations by Karamitros D. et al. [47], with the vast majority belonging to the “common myeloid progenitor” population (88%), followed by the megakaryocyte/erythroid progenitor compartment (6%). This suggested that sub-cluster 3 cells might consist of multipotent progenitors which, in contrast to HSC, are known to be highly proliferative and metabolically active [49, 50].

Within our population, half of the 115 abundantly/differentially expressed genes were almost exclusively associated with sub-cluster 3, followed by shared expression with promyelocytes, erythrocytes/megakaryocytes, activated neutrophils, and monocytes (S7 Dataset). Thirty-six of these genes were also present in CMV^+^ cells, with the majority being shared with the promyelocytes and erythrocytes/megakaryocytes clusters. Together, these data indicate that sub-cluster 3 is comprised of proliferating cells expressing erythroid, monocytic and granulocytic markers, which we surmised might represent CFU-GEMM oligopotent progenitors.

To further test this hypothesis, cells belonging to the cluster 7, erythro, mono, MDDC, CMV^+^, promyelo, act neut and sub-cluster 3 groups depicted in Fig 4C were ordered along trajectories corresponding to their inferred differentiation pathways using Monocle [51]. A trajectory with four main branches extending from a rooted center was generated (Fig 5A), and the identity of cells composing each of the eight groups was uncovered using Seurat [52] (Fig 5B and S8 Fig). Cells in group D, the root center, expressed the same key genes as sub-cluster 3 cells in Fig 4C, while its closest neighbors, group E, F, G and H, expressed markers typical of the monocytes, erythrocytes, promyelocytes and activated neutrophils clusters in Fig 4C, respectively (S8 Fig and S9 Dataset). The most isolated cluster of cells, group B, was related to CL7 in Fig 4C, and differentially expressed CD52 and FCER1A (S9 Dataset). Thus, the observed pseudotime distances and cluster organization strongly implicated group D as the most likely origin of erythrocytes, megakaryocytes, promyelocytes, neutrophils and monocytes, suggesting that cells comprising group D/sub-cluster 3 might indeed represent CFU-GEMM. As the CMV^+^ cluster was immediately adjacent to group D, we further conclude that CMV^+^ cells are directly related to, and possibly derived from, CFU-GEMM progenitors.

**Figure 5.**
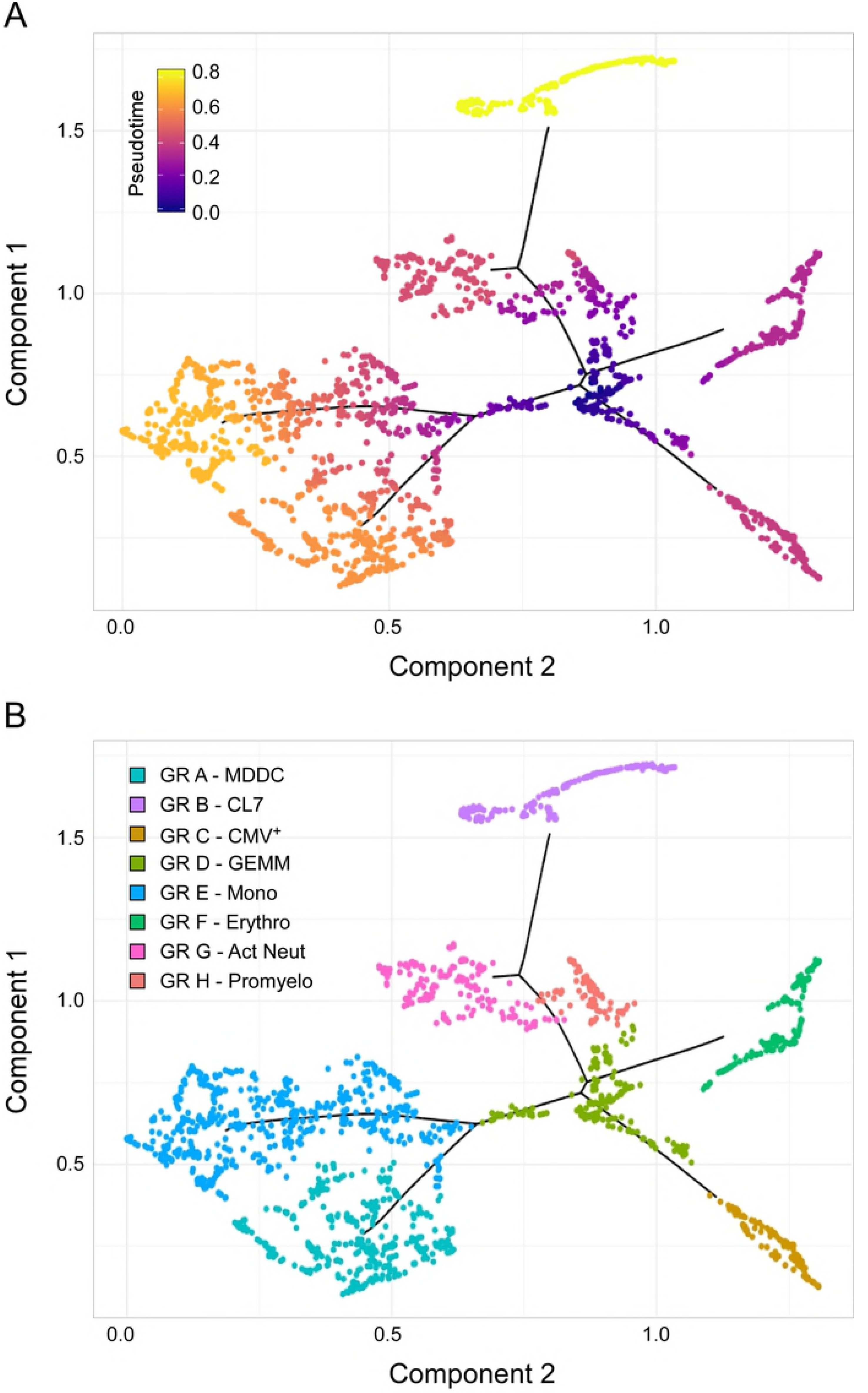
Identification of sub-cluster 3 as the origin of promyelocytes, activated neutrophils, erythrocytes, megakaryocytes, monocytes, and CMV^+^ cells. (A) Pseudotime ordering of data from cells belonging to the CL7, erythro, mono, MDDC, CMV, promyelo, act neut and sub-cluster 3 groups shown in Fig 4C into a two-dimensional component space using Monocle. The main path of the minimum spanning tree is depicted by solid black lines arising from a central root of cells with a pseudotime of zero (dark blue dots), and branching outward to clusters with higher pseudotime values, representing differentiated cell types (purple, orange and yellow dots). (B) Cell group labeling based on the expression of key marker genes identified with Seurat. GR = group; MDDC = monocyte-derived dendritic cells; CL7 = CL7 from Fig 4C; GEMM = colony-forming unit-granulocyte, erythrocyte, monocyte/macrophage, megakaryocyte; Mono = monocytes; Erythro = erythrocytes; Act Neut = activated neutrophils; Promyelo = promyelocytes.

### The majority of the genes characterizing CFU-GEMM cells are more highly expressed in this cluster than in the rest of the population, and are maintained to similar levels in CMV^+^ cells

To understand why cells in sub-cluster 3 (relabeled GEMM) in particular allowed infection to initiate we sought to identify which set of genes and, consequently, which cellular functions, were most differentially regulated in GEMM and CMV^+^ cells with respect to the rest of the population. A set of 1989 genes was identified by the 10X Genomics Cell Ranger software as being more selectively expressed in the CMV^+^ and GEMM clusters relative to all other clusters. The majority of these genes (1361, or 68% for GEMM, and 1460, or 73% for CMV^+^ cells) were associated with positive Log_2_ fold change values, indicating that most of the GEMM-specific genes were more highly expressed in these cells than elsewhere, and that, as expected, infection was accompanied by a strong transcriptional up-regulation of cellular genes (S10 Dataset, sheet 1).

The differentially expressed genes were then partitioned between “synchronous” and “asynchronous”, depending on whether their transcription was similarly regulated in GEMM and CMV^+^ cells or not. Genes that were up-regulated in GEMM cells relative to the rest of the population, and that were expressed to similar levels or further up-regulated in CMV^+^ cells (total = 1077), as well as genes that were down-regulated in GEMM cells and that were expressed to similar levels or further down-regulated in CMV^+^ cells (total = 325) were considered synchronous, while genes that were up-regulated in GEMM but down-regulated at least two-fold in CMV^+^ cells, and vice-versa, were labeled asynchronous (total = 587). The majority (1402, 70 %) of the selected genes fell into the synchronous category. Of these, most were up-regulated in the GEMM cluster and expressed to similar levels in CMV^+^ cells, with only 28 genes being further induced in infected cells, suggesting that GEMM cells already contain large numbers of transcripts beneficial (or neutral) to infection (S10 Dataset, sheet 1). As levels of down-regulated genes were also mostly maintained without any further repression by infection, we hypothesized that GEMM cells might be preferentially infected because their transcriptional landscape requires the least amount of optimization by viral effectors.

### Expression of genes with functions in energy production, cell cycle control, RNA and protein metabolism is higher in both GEMM and CMV^+^ cells

To pinpoint the functional areas distinguishing GEMM cells from the remainder of the population, the most differentially expressed synchronous genes, and all of the asynchronous genes (1304 in total) were partitioned into 15 categories based on their encoded functions (S10 Dataset, sheet 2). The transcript abundance of each gene found in the CMV^+^ or GEMM clusters was then divided by the abundance in the rest of the cells (CMV/REST and GEMM/REST) or in GEMM cells (CMV/GEMM), and the distributions of the Log_2_ ratio values were plotted (Fig 6). As expected, only the GEMM/REST and CMV/REST, but not the CMV/GEMM ratio distributions of all genes were identified by the Wilcoxon signed rank test as having Log_2_ median values significantly different from zero (Fig 6A). Genes with roles in mitochondrial functions (Fig 6I), proliferation and cell cycle control (Fig 6J), RNA metabolism (Fig 6L) and protein processing (Fig 6K) were also more highly expressed in both GEMM and CMV^+^ cells, and were thus further scrutinized.

**Figure 6.**
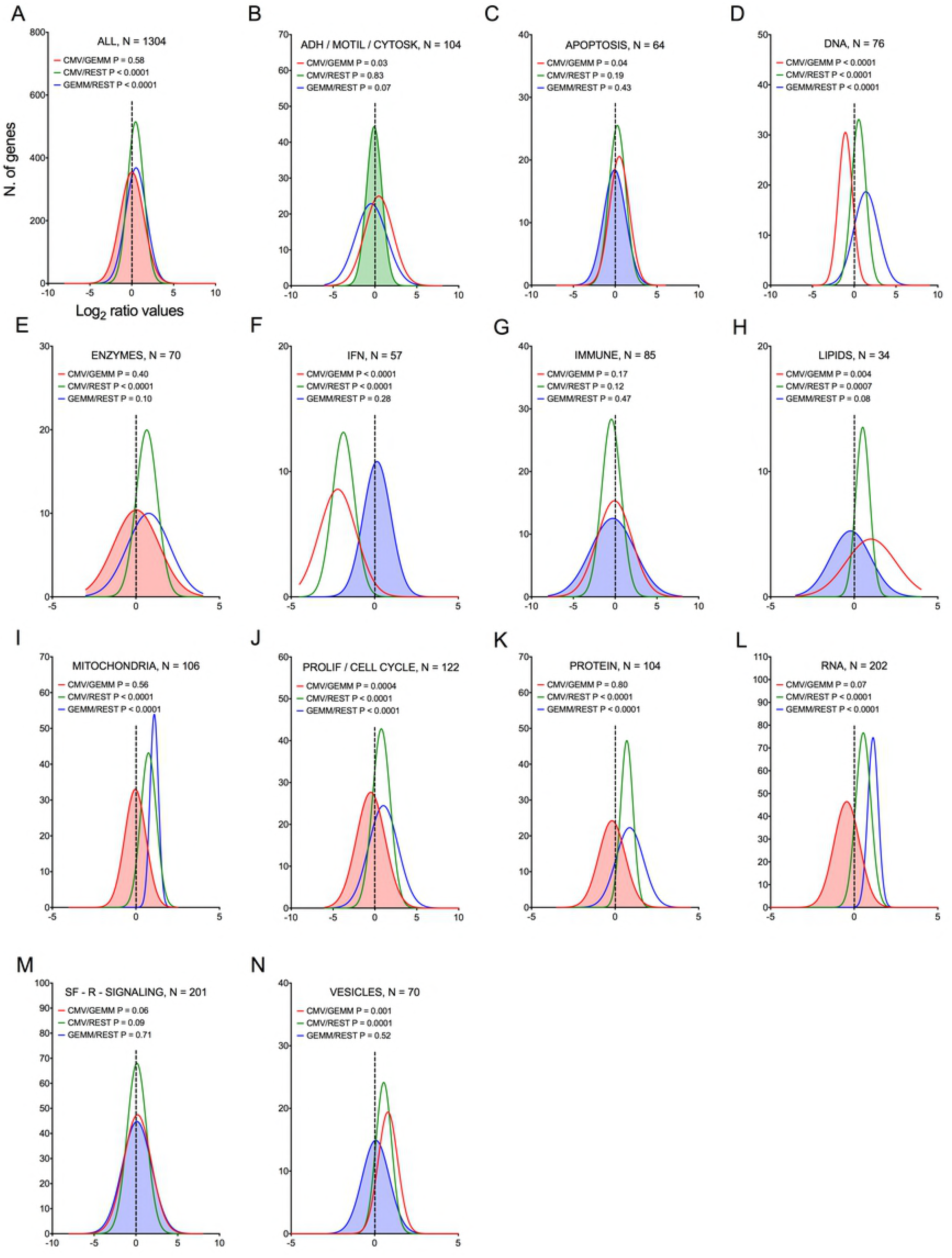
Differential expression of genes belonging to multiple functional categories in GEMM cells, CMV^+^ cells, and in the rest of the population. Log_2_ ratio value distributions obtained by dividing the mean number of transcripts/cell for each gene in the CMV^+^ or GEMM clusters by the mean number of transcripts/cell in the rest of the cells (CMV/REST, green line, and GEMM/REST, blue line) or in GEMM cells (CMV/GEMM, red line). The distribution obtained from all genes is shown in (A), while B-N show the distributions of genes falling in each functional category. The Wilcoxon signed rank test was used to identify populations with median values significantly different from zero. The population with the lowest P value is highlighted by coloring of the area under the curve. The dashed line marks the ratio = 1 point. N = number of genes in each category; ADH/MOTIL/CYTOSK = adhesion/motility/cytoskeleton; IFN = interferon; PROLIF/CELL CYCLE = proliferation/cell cycle; SF-R-SIGNALING = soluble factors/receptors/signaling.

#### Mitochondria

Genes involved in ATP production, mitochondrial protein synthesis, and mitochondrial transport were consistently more abundant in GEMM cells than elsewhere (Fig 7A and C-E, blue lines), with their expression levels remaining largely unchanged in CMV^+^ cells (Fig 7A, and C-E, red lines). Among these, genes encoding members of the ATP synthase and NADH dehydrogenase complexes of the electron transfer chain were the most represented, together with genes encoding mitochondrial ribosomal proteins (S10 Dataset, sheet 3). This suggests that infection might preferentially start in GEMM cells due to the existence of an intracellular environment already geared toward high energy production, and hence capable of supporting the large metabolic requirements of viral replication. We did indeed previously observe a similarly strong up-regulation of genes with functions in oxidative phosphorylation and fatty acid β-oxidation in infected fibroblasts at late times pi [53], indicating that the enhancement of mitochondrial functions is a key feature of infection.

**Figure 7.**
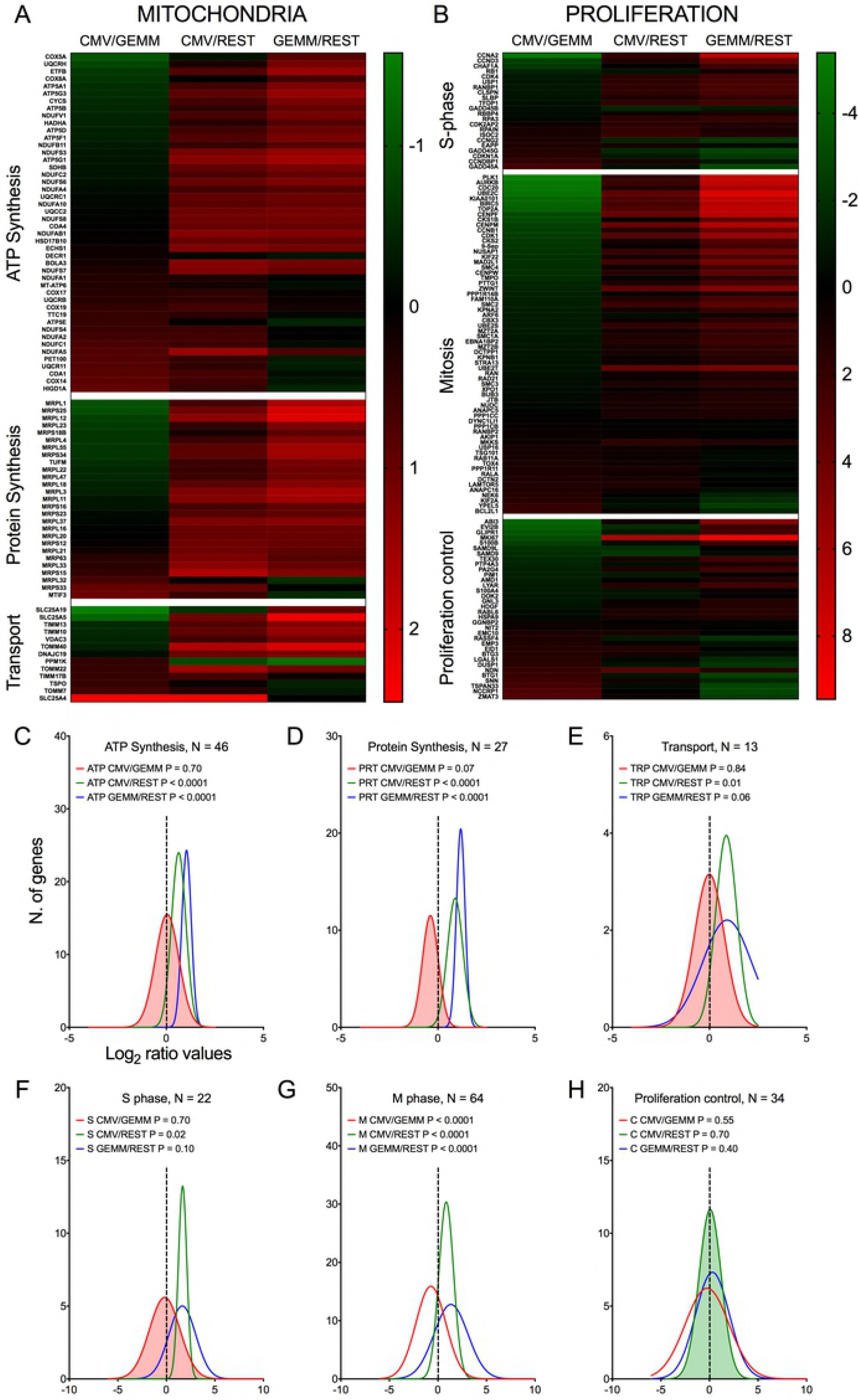
Differential expression of genes with roles in mitochondrial function and proliferation control in GEMM cells, CMV^+^ cells, and the rest of the population. Heatmap (A and B) and distributions (C-H) of Log_2_ ratio values obtained by dividing the mean number of transcripts/cell of genes with roles in mitochondrial functions (A and C-E) and in proliferation control (B and F-H) as found in the CMV^+^ or GEMM clusters by the mean number of transcripts/cell in the rest of the cells (CMV/REST, green line, and GEMM/REST, blue line) or in GEMM cells (CMV/GEMM, red line). The heatmap color scales refer to the Log_2_ ratio values. The Wilcoxon signed rank test was used to identify populations with median values significantly different from zero. The population with the lowest P value is highlighted by coloring of the area under the curve. The dashed line marks the ratio = 1 point. N = number of genes in each category.

#### Proliferation/cell cycle

Consistent with the notion that multipotent progenitors are highly proliferative [49, 50], GEMM cells expressed higher levels of genes encoding S and M phase effectors than the rest of the population (Fig 7B and F-G, blue lines and S10 Dataset, sheet 4). CMV infection of fibroblasts was reported by us and others to repress expression of genes promoting entry into S phase, while simultaneously inducing expression of DNA synthesis effectors [53–56]. In keeping with these observations, CMV^+^ cells contained lower transcript amounts of genes promoting entry into S phase, such as CCNA2, CCND3, MKI67 and RB1, but higher transcript levels of genes encoding inhibitors of S phase progression, including BTG1, BTG3, CCNDBP1, CDKN1A (Cip1) and the HSC quiescence-promoting gene NDN [57], which was almost exclusively expressed in CMV^+^ cells (not shown). Transcription of DNA replication effectors was, by contrast, inconsistently induced. While expression of some genes, such as the catalytic subunit of the DNA polymerase delta (POLD2) and its interacting protein POLDIP2, RPA3 and the RPA complex nuclear importer RPAIN, was high, transcription of others such as PCNA, MCM3, MCM7, and FEN1 was reduced in CMV^+^ cells. We speculate that this mixed transcriptional regulation might be typical of the early phase of infection, when viral factors are still in the process of gaining control over cell proliferation, while at later times, when data from fibroblasts were collected [53], viral DNA synthesis is already fully established.

We previously reported that CMV infection induces the appearance of aberrant mitotic figures, supported by the induction of numerous genes involved in M phase progression [53]. Although this feature was shared by different CMV strains, it was by far most evident with the attenuated strain AD169 than with TB40/E [58]. Consistent with the TB40/E pattern, only a minority of the 63 genes with functions in mitosis were maintained to high levels in CMV^+^ cells, while the rest were down-regulated (Fig 7B and G), including the two main components of the mitosis-promoting factor, CDK1 and CCNB1, chromatin condensation agents (SMC2, SMC4, ZWINT and TOP2A), mitotic spindle assembly controllers (AURKB, BIRC5, PLK1, MAD2L1, and CENPF), components of the anaphase-promoting complex (CDC20 and PTTG1), and cytokinesis effectors (SEPT9, ARF6, and RAB11A).

Together, these data are consistent with a CMV-induced block in cell proliferation, aimed at curtailing usage of cellular resources for processes irrelevant to viral replication, such as mitosis, and steering others, such as those devoted to cellular genome replication, toward viral DNA production instead.

#### RNA metabolism

As expected for metabolically active cells, expression of numerous genes involved in RNA processing, splicing and translation were more highly expressed in GEMM and in CMV^+^ cells than in the rest of the population (Fig 8A, C and D, blue and green lines and S10 Dataset, sheet 5). By contrast, expression of ∼ 70% of transcription-related genes was similar in GEMM and in the rest of the cells, but was up-regulated in CMV^+^ cells (Fig 8A and B, blue and red lines).

**Figure 8.**
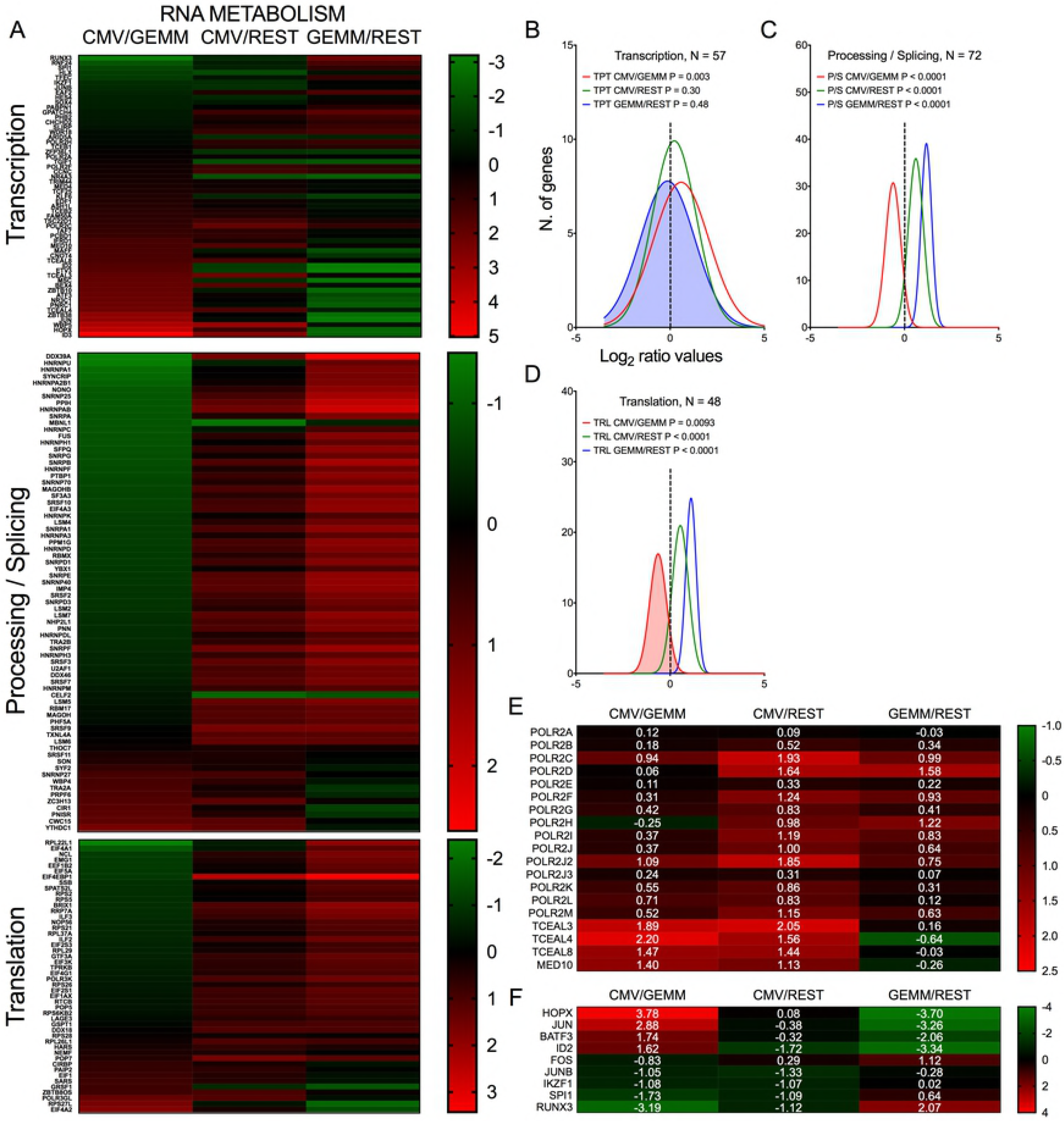
Differential expression of genes with roles in RNA metabolism in GEMM cells, CMV^+^ cells, and the rest of the population. Heatmap (A, E and F) and distributions (B-D) of Log_2_ ratio values obtained by dividing the mean number of transcripts/cell of genes with roles in RNA transcription, processing and translation as found in the CMV^+^ or GEMM clusters by the mean number of transcripts/cell in the rest of the cells (CMV/REST, green line, and GEMM/REST, blue line) or in GEMM cells (CMV/GEMM, red line). The heatmap color scales refer to the Log_2_ ratio values. Numbers in white font in E and F report the Log_2_ ratio values of each gene. The Wilcoxon signed rank test was used to identify populations with median values significantly different from zero. The population with the lowest P value is highlighted by coloring of the area under the curve. The dashed line marks the ratio = 1 point. N = number of genes in each category.

Particularly revealing of the strong impetus of infection toward stimulating cellular gene transcription on a broad scale was the induction of several RNA polymerase II subunits and elongation factors (Fig 8E), while among transcription factors, the most strongly up-regulated in CMV^+^ cells were the HOPX homeobox (CMV/GEMM ratio ∼14-fold), the proto-oncogene JUN (7-fold) with its heterodimerization partner BATF3 (3-fold), and the differentiation inhibitor ID2 (3-fold). Transcription of the other two JUN partners, FOS and JUNB, and of the regulators of hematopoietic cells differentiation IKZF1, SPI1/PU.1, and RUNX3 was instead reduced 2- to 9-fold (Fig 8F).

Thus, the early stages of infection appear to be associated with a sharp push towards increased production of RNA synthesis and processing effectors. This is likely required to support viral gene transcription in order to fine-tune viral control over a variety of cellular processes, including cell differentiation.

#### Protein metabolism

In keeping with the robust infection-associated stimulation of gene translation, expression of numerous protein chaperones and post-translational modifiers was also higher in both GEMM and CMV^+^ cells than in the rest of the population (Fig 9A and B-C, blue and green lines and S10 Dataset, sheet 6). Chaperone-assisted protein folding occurs via three main routes, the simplest one being via interactions with single HSP70 or HSP90 family members. Some polypeptides require the sequential binding of HSP70 and HPSP90 instead, while others need the intervention of the chaperonin containing TCP1 complex (CCT) [59]. Both HSP70 coding transcripts, HSPA1A and HSPA1B, and their co-chaperone DNAJB6 were expressed to lower levels in GEMM cells than in the rest of the population, and were up-regulated in infected cells. The adaptor protein STIP1, which coordinates protein transfer from HSP70 to HSP90, the inducible (HSP90AA1) and constitutive (HSP90AB1) HSP90 isoforms, and all eight subunits of the CCT complex were expressed at higher levels in both GEMM and CMV^+^ cells. A similar pattern of regulation was observed for calnexin (CANX) and calreticulin (CALR), and for seven out of eleven members of the large endoplasmic reticulum (ER)-localized multiprotein complex (HSPA5, DNAJB11, HSP90B1, PPIB, PDIA6, SDF2L1 and ERP29), which, together, comprise the ER protein quality control system [60, 61] (Fig 9E).

**Figure 9.**
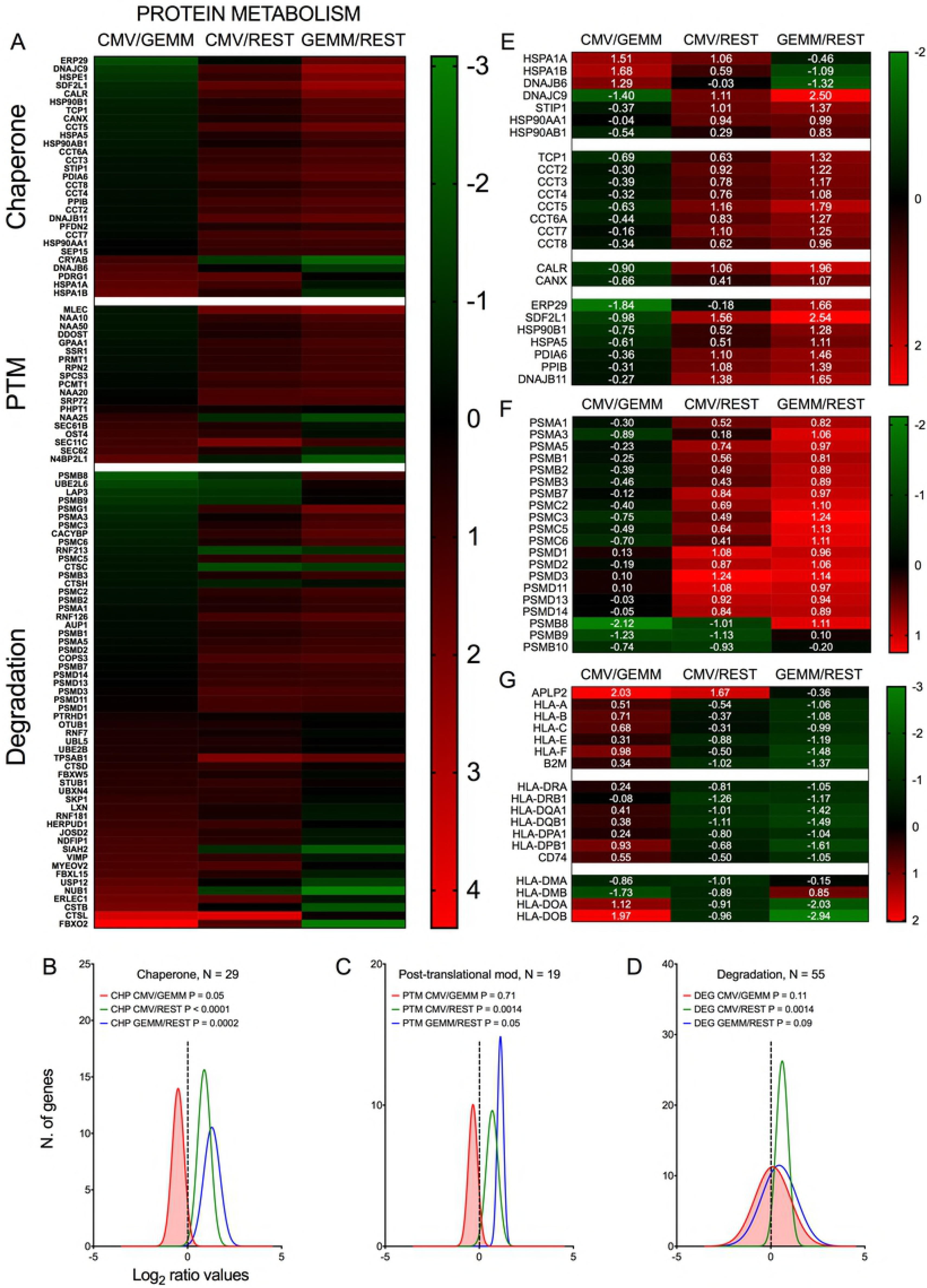
Differential expression of genes with roles in protein metabolism and antigen presentation in GEMM cells, CMV^+^ cells, and the rest of the population. Heatmap (A, E, F and G) and distributions (B-D) of Log_2_ ratio values obtained by dividing the mean number of transcripts/cell of genes with roles in protein metabolism as found in the CMV^+^ or GEMM clusters by the mean number of transcripts/cell in the rest of the cells (CMV/REST, green line, and GEMM/REST, blue line) or in GEMM cells (CMV/GEMM, red line). The heatmap color scales refer to the Log_2_ ratio values. Numbers in white font in E-G report the Log_2_ ratio values of each gene. The Wilcoxon signed rank test was used to identify populations with median values significantly different from zero. The population with the lowest P value is highlighted by coloring of the area under the curve. The dashed line marks the ratio = 1 point. N = number of genes in each category.

Levels of numerous genes with roles in protein degradation were also slightly higher in GEMM and CMV^+^ cells (Fig 9A and D, blue and green lines). Of particular interest was the up-regulation of 17 subunits (out of 33) of the proteasome (Fig 9F). Protein degradation may benefit the virus by removing unwanted cellular polypeptides and damaged or misfolded proteins, while simultaneously enhancing amino acid availability. An essential role of the proteasome, however, is to produce antigenic peptides suitable for presentation on major histocompatibility complex (MHC) class I molecules, an activity extremely detrimental to virus spread. In the immunoproteasome, the proteolytic subunits PSMB5, 6 and 7 are replaced with PSMB8, 9 and 10. Very intriguingly, and consistent with data from infected fibroblasts [62], expression of these latter subunits was down-regulated in CMV^+^ cells (Fig 9F).

In addition to curtailing the ability of the immunoproteasome to produce antigenic peptides, MHC class I activities were also negatively impacted by the strong transcriptional induction of APLP2, an enhancer of MHC class I internalization and turnover [63, 64]. Rather intriguingly, transcript levels of genes encoding the three main MHC class I molecules, HLA-A, -B, and -C, and of their binding partner B2M, as well as of the three main MHC class II isotypes, HLA-DR, -DQ, and -DP and the invariant chain CD74 were already ∼ 2.5-fold lower in GEMM cells than in the rest of the population and were not further reduced in CMV^+^ cells (Fig 9G). By contrast, expression of HLA-DMA and HLA-DMB, which assist in the binding of high affinity antigenic peptides into MHC class II [65], were repressed while transcription of HLA-DOA and HLA-DOB, which increase tolerance to self-peptides [65], was increased (Fig 9G). Together, these data underscore the strong effects of infection on fine-tuning the cellular protein “portfolio” to match the virus’ needs, and highlight the pristine selectivity of viral effectors in modulating the expression of specific cellular proteins in order to protect infected cells from detection and elimination by the host immune system.

### Expression of genes with functions in IFN-mediated antiviral defenses is similar in GEMM and in the rest of the cells, and is strongly down-regulated in CMV^+^ cells

Akin to genes belonging to categories of apoptosis, immune, lipids, soluble factors/receptors/signaling and vesicles (Fig 6C, G, H, M and N, blue line), transcript levels of IFN-related genes were overall similar in GEMM cells and in the rest of the population (Fig 6F, blue line). Very excitingly, however, this category contained the most strongly down-regulated genes of all in CMV^+^ cells (median Log_2_ CMV/GEMM ratio value of – 1.9, P < 0.0001, Fig 6F, red line).

Compared to the rest of the population, GEMM cells contained higher levels (median ratio, ∼ 1.5-fold) of transcripts encoding sensors of viral double-stranded DNA and RNA, such as IFI16 [66], HMGB1 [67], DDX58/RIG-I [68], IFIH1/MDA5 [69] and EIF2AK2/PKR [70], of signaling mediators like STAT1, and of transcriptional activators such as IRF3, IRF7 and IRF8 [71], but lower levels (median ratio, ∼ 4-fold) of negative regulators of IFN production and signaling such as IRF2 [72], IRF4 [73], TRAFD1 [74] and SOCS1 [75]. Expression of IFN effectors including IFIT1, IFIT2 and IFIT3, which recognize and prevent translation of virally produced triphosphorylated RNA molecules [76], IFITM1, IFITM2 and IFITM3, which block infection at multiple steps including entry [77], ISG15 and its conjugating (HERC5) and de-conjugating (USP18) enzymes, which disrupt the activity of viral proteins by ISGylation [78], as well as of known (MX1, MX2, OAS1, OAS2, OAS3 and OASL) [79, 80], or suspected anti-viral proteins such as viperin [81], SAMHD1 [82] and ISG20 [83] were instead similarly abundant in GEMM and the rest of the cells (median ratio, ∼ 1.1-fold) (Fig 10, GEMM/REST column).

**Figure 10.**
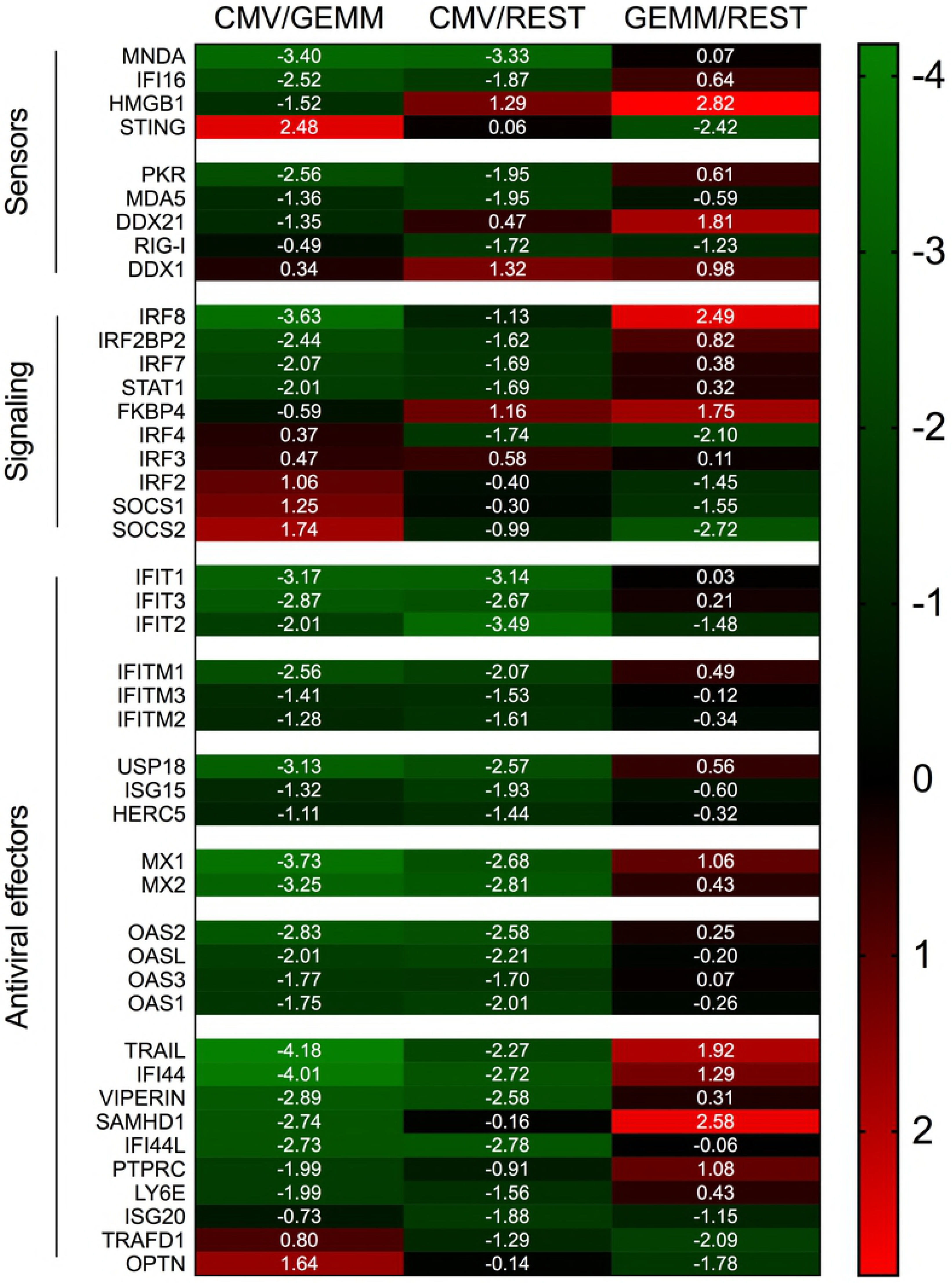
Differential expression of IFN-related genes in GEMM cells, CMV^+^ cells, and the rest of the population. Heatmap of Log_2_ ratio values obtained by dividing the mean number of transcripts/cell of IFN-related genes as found in the CMV^+^ or GEMM clusters by the mean number of transcripts/cell in the rest of the population (CMV/REST and GEMM/REST) or in GEMM cells (CMV/GEMM). The heatmap color scale refers to the Log_2_ ratio values. Numbers in white font report the Log_2_ ratio values of each gene.

Together, these data suggest that GEMM cells are not defective in their ability to detect, respond and potentially antagonize viral infection. Rather, GEMM cells appear to be similarly, or even more responsive than the rest of the population, indicating that the lack of appropriate cellular defenses is unlikely to be the main reason for their preferential infection. Interestingly, and similar to the situation with MHC class I and II genes, transcriptional modulation of these genes in CMV^+^ cells appeared to be selective: while mRNA levels of most IFN antiviral effectors were powerfully reduced, transcription of negative regulators was enhanced, with the notable exception of the adaptor protein TMEM173/STING [84–87] and the TBK1 activator OPTN [88], which are both involved in IFN production following CMV DNA detection by MB21D1/cGAS [89]. Taken together, these findings provide support to our theory whereby infection preferentially begins in GEMM cells due to their higher metabolic, proliferative, and RNA and protein synthesis rates, rather than to impairments in their capacity to mount strong cellular defenses.

## DISCUSSION

In previous studies, we showed that activated and non-activated myeloid cells differentiated from CD34^+^ HSC are semi-permissive to CMV infection [9, 14–16]. As such, they constitute a useful model to identify the cellular determinants of viral tropism.

Based on our current and previous data, resistance to infection appears to be multilayered, affecting multiple sequential steps in the viral life cycle, and progressively narrowing the proportion of cells capable of supporting the full viral replication cycle. While in homogeneous permissive cell populations such as fibroblasts the probability for a cell to remain free of viral particles at an MOI of ten is null, our data show that at day one pi ∼ 40% of activated myeloid cells do not contain any viral RNAs (Fig 2). However, of the CMV-transcript^+^ cells, only ∼ 7% express the UL122 and UL123 ORFs necessary for infection onset, and of these, about half contain additional transcripts needed for CMV genome replication. While the proportion of cells progressing toward the replicative stage may increase over time, the number of IE1/IE2^+^ cells remained unchanged from day two pi onwards (Fig 1), suggesting that the cellular barriers restricting infection onset are never overcome. Despite the presence of donor-dependent variation (expected not only for primary cells but especially for hematopoietic cell types), activated cells consistently contained larger proportions of IE1/IE2^+^ cells but produced less progeny (Fig 1). We used these less permissive cells as a tool to identify the cellular pathways involved in inhibiting (or promoting) infection.

Although the absence of viral RNAs in ∼ 40% of activated cells may depend, at least in part, on timing and detection limits, it is also possible for this portion of the population to be more resistant to viral entry due to the presence of specific restriction factors and/or the absence of entry facilitators. However, no specific cellular genes were identified as being selectively transcribed in viral RNA^+^ or RNA^−^ cells. Transcripts coding for proteins currently known to support virion entry were also either absent (EGFR, THY1/CD90 and ITGB3) or found in only a minute proportion of cells (ITGAV, ITGA2, ITGA6 and PDGFRA), while more abundant levels of transcripts coding for BSG and ITGB1 did not selectively partition with viral RNA^+^ cells. Because our myeloid cell cultures are highly heterogeneous (Fig 4), preferential infection of select sub-groups may still have been facilitated by the expression of specific genes. BSG, for instance, was present in 99% of GEMM cells but in only 60% of cluster 7 cells. Conversely, viral RNA*−* cells may have resisted infection owing to the expression of subset-specific molecules. However, the fact that no “universal” entry resistance/enabling gene(s), expressed by all viral transcript^+/−^ cells, could be identified implies that such gene(s) may not exist. This is in contrast to other cell types such as endothelial and epithelial cells, whose infection instead depends on the expression of surface molecules (such as BSG), acting as receptors for specific glycoprotein complexes present on the virion’s surface [37].

Only a small fraction (∼ 3%) of CMV-transcript^+^ cells expressed multiple viral ORFs at high levels. Presumably, these represent cells that will progress toward lytic replication, as corroborated by the presence of early viral proteins in a similar proportion of cells (Fig 1 and Fig 3). Thus, we wondered about the fate of infection in the remainder of the cells, which contain lower amounts of viral transcripts, and no detectable viral proteins. Intriguingly, a similar scenario was recently encountered following single-cell RNA sequencing of TB40/E-infected CD14+ and CD34^+^ cells [90]. Elevated levels of viral transcripts were observed in just ∼ 2% of monocytes, while the rest of the population, which contained lower amounts of a wide range of viral transcripts, were interpreted as potentially being latently infected. This led us to speculate that the remaining CMV-transcript^+^ cells in our population might be either latently infected or on a path toward latency. While this hypothesis requires additional testing, it remains a thrilling possibility, especially in view of recently presented evidence supporting the potential association of viral latency with quantitative rather than qualitative changes in viral gene expression [90]. Alternatively, it is of course possible for viral transcripts to simply be detected and eliminated by cellular defense mechanisms, producing an abortive infection.

Although expression of CD207/langerin and CD1a was observed in a number of cells within the population, Langerhans cells did not appear to be the main source of infected cells, at least at day one pi. Rather, multiple lines of evidence indicate that CMV^+^ cells derive from a cluster with the hallmarks of GEMM colony forming units, albeit devoid of transcripts coding for some of the markers traditionally used to describe this population, *i.e.* CD34, CD38, and CD123. As none of the genes we and others [47, 48] found to be selectively expressed by these cells encode surface molecules, their isolation from either *in vitro* differentiated myeloid populations or hematopoietic tissues is particularly challenging. Consequently, we do not currently have direct evidence that this specific cell type can support CMV lytic infection *in vivo*.

Very recent data from single-cell RNA-seq analyses of hematopoietic processes have revealed that lineage development is a continuous process, more usefully depicted by Waddington’s landscapes [91], than by more rigid cell differentiation trees. In this emergent scenario, CD34^+^ HSC are visualized as beads rolling along a surface stretching from a higher to a lower point in space, and containing ridges and valleys. These ridges, corresponding to barriers separating individual lineages, are smaller near the top and become increasingly higher towards the bottom as expression of fate mediators progresses in each cell. Once ridges become too high cells can no longer change their identity and terminal lineages are established [47, 48, 92, 93]. We believe that the permissive cell type we identified in this study corresponds to a mid-point along this surface, characterized by the loss of pluripotency, but not yet enclosed by the high ridges separating granulocytes, monocytes, erythrocytes and megakaryocytes from each other. While this cell type is likely to exist *in vivo*, it may have been missed in previous studies of CMV tropism due to its rarity, and/or to the lack of specific surface markers.

An interesting question in this regard is: when did these cells arise during CD34^+^ HSC differentiation, and what factors influence this process? The CD34^+^ cells we employed in this study were isolated from cord blood and were amplified for 8-10 days in the presence of FL, stem cell factor (SCF), and thrombopoietin (TPO) before differentiation. Others have shown that GEMM cell numbers increase by 850-fold during culture of CD34^+^ cord blood cells in the presence of FL and TPO for 15 weeks [94], while CD34^+^ HSC from peripheral blood produce lower total cell numbers and colony forming units than CD34^+^ HSC from cord blood after three weeks of exposure to FL plus TPO or FL plus TPO plus SCF [95]. Rather interestingly, HSC expansion was also associated with the rapid loss of the CD34 marker [95]. These data suggest that cord blood-derived HSC might have a stronger propensity to develop into GEMM cell-containing populations than cell populations isolated from peripheral blood or bone marrow. FL, SCF, and TPO promote self-renewal of CD34^+^ cells and have been used to expand cord blood HSC *in vitro* for therapeutic intervention [94–98]. While all three cytokines stimulate HSC division, TPO also drives megakaryocyte development [99], and FL, which steers hematopoiesis toward the lympho-myeloid lineage at the expense of erythrocytes/megakaryocytes, is essential for the generation of dendritic cells [100]. Thus, in addition to stimulating HSC proliferation, these cytokines may have provided the very first “ridges”, nudging HSC differentiation toward GEMM cells. Intriguingly, we were able to detect the presence of progeny virus in the culture supernatant of amplified (but not of non-amplified) CD34^+^ cells exposed to TB40/E, albeit with low frequencies (not shown). This led us to wonder if, perhaps, GEMM cells might be present in amplified HSC cultures even before exposure to the differentiation cocktail. Thus, cell culture conditions are critical when studying HSC in conjunction with CMV. CD34^+^ cells are extremely plastic, and can clearly give rise to clustered sub-populations of myeloid cells, some of which permissive to lytic infection. Being a minority in the population these clusters can easily escape detection and may introduce unwanted and unnoticed “lytic noise” [90] in studies of viral latency.

While the reason for the preferential infection of CMV-transcript^+^ cells remains unclear, our data provide a plausible rationale for initiation of lytic infection in GEMM cells; *i.e.*, their higher expression of multiple gene products involved in energy, RNA, and protein production, as well as in cell cycle control, which likely create an intracellular environment particularly conducive to infection onset by lowering the amount of energy required from viral effectors to steer cellular processes away from cell needs and toward viral replication.

We initially reported the up-regulation of numerous genes with functions in mitochondrial oxidative phosphorylation, fatty acid β-oxidation, and malate-aspartate, ATP, and citrate transport systems in infected fibroblasts at late times pi [53]. Our findings were subsequently confirmed and expanded by a number of studies in fibroblasts and other cell types [101–104]. To our knowledge, the current work is the first report showing that a strong transcriptional induction of this type of genes also occurs in myeloid cells and at early times post-entry (Fig 7), making it a hallmark of CMV infection in different cell types and a requirement for successful viral replication.

The cell cycle is also a very well-known target during viral infection. Multiple studies have shown that CMV infection drives host cells into the G1/S phase to shunt cellular resources required for DNA synthesis and repair toward viral rather than cellular genome replication. Manipulation of cell cycle functions occurs at multiple levels, including protein transcription, translation, stability, posttranslational modification, and subcellular localization [105]. We previously showed that CMV infection is associated with a very strong positive impact on the expression of multiple S phase, M phase, and DNA activity regulators in fibroblasts, leading to the appearance of aberrant mitotic figures, which we called pseudomitosis, at late times pi [53, 58]. Here, we found that expression of genes involved in S phase control was higher in GEMM cells and remained high in CMV^+^ cells, whereas transcription of M phase regulators was reduced (Fig 7). While fibroblasts were infected at confluency (when the majority of the cells are in G0/G1), GEMM cells were likely actively proliferating at the moment of contact with CMV. We thus believe that our new data highlight the exquisite ability of infection to fine tune its impact on gene transcription according to the conditions of the cell at the time of entry, in order to reach optimal expression levels of specific genes useful to viral replication. Although entry into mitosis is clearly detrimental to viral replication [106], the presence of select M phase proteins may be needed to perform specific tasks, such as viral genome compaction, disentangling, or transport. To reach an ideal protein concentration these genes may thus need to be transcriptionally upregulated in quiescent fibroblasts, whereas downregulation may prevail when cells are already actively cycling.

High expression levels of genes involved in RNA processing, splicing, and translation were not unexpected for metabolically active cells such as GEMM, and neither was it surprising that they were maintained in CMV^+^ cells (Fig 8). Indeed, a similar scenario was observed by us [53] and others [107] in infected fibroblasts. Again, our data validate and expand these findings to include myeloid cells, clearly marking these metabolic processes as pivotal for successful infection.

Expression of several genes with essential roles in hematopoietic development was also altered in CMV^+^ cells (Fig 8). These include HOPX, which regulates primitive hematopoiesis [108], BATF3, vital for the development of conventional cross-presenting CD8α^+^ dendritic cells, ID2, whose expression in CD34^+^ HSC inhibits the development of dendritic cell precursors [109], RUNX3, whose depletion leads to defects in the proliferation and differentiation of activated cytotoxic CD8+ T cells, helper Th1 cells and NK cells, and to the disappearance of skin Langerhans cells [110], IKZF1, essential for normal lymphopoiesis and for myeloid, megakaryocyte and erythroid differentiation [111], and SPI1/PU.1, which is critical for the generation of all hematopoietic lineages [112]. If dysregulated expression of these genes also occurs in infected progenitors *in vivo*, it may powerfully affect the development and functions of multiple arms of the hematopoietic system, providing potential new culprits for the infection-associated problems ensuing congenital infection and hematopoietic stem cell transplantation.

Finally, we observed a much stronger negative impact of infection on the expression of genes with functions in the production and responses to IFN than previously reported, accompanied by the induction of a very small, and possibly selected, number of genes. To our knowledge this is the first analysis of transcriptional responses to CMV infection in myeloid cells conducted at the single-cell level and, hence, capable of comparing gene expression levels in lytically infected cells to those in bystander cells co-existing within the same population. Our data thus provide a new perspective on how host defenses are raised and subsequently offset by the virus, in contrast to previous analyses that compared mean gene expression levels in CMV-infected samples to those in separate mock-infected cells [113–117]. Aside from the detection method, differences may also depend on the time pi, the cell type, and the strain of virus used. Infection recognition was shown to occur very rapidly in monocytes and fibroblasts, leading to the activation of the transcription factors IRF3 and NF-κB, and to the implementation of the IFN transcriptional program within 4-8 hours [113, 114, 116, 118–122]. Structural components of the virion, such as the tegument proteins pp65 and/or pp71 [113, 114, 123, 124], as well as viral immediate-early and early proteins [125, 126] then cooperate to blunt these responses via multiple mechanisms, including inactivation of the double-stranded DNA sensor MB21D1/cGAS, blockage of the STING-TBK1-IRF3 complex assembly, inhibition of NF-kB binding to DNA, and degradation of the signal transduction molecules JAK1 and STAT2 [123, 124, 127–132]. IFN-related genes that are highly transcribed at 4-8 hours pi may thus be downregulated at 24 hours pi, or upon full implementation of viral countermeasures. The importance of viral anti-IFN defenses is indeed underscored by the fact that five of the nine genes more abundantly expressed in CMV^−^ cells encode IFN-induced antiviral proteins (MX1, OAS1, OAS2, IFIT3, and USP18, Supplementary Fig 1), suggesting that effective downregulation of their expression could not be achieved in the absence of specific viral gene products.

Basal expression of sensors, signal transducers, and IFN-inducible genes as well as the speed and strength whereby antiviral responses are mounted can also be cell type dependent [133]. IRF3 and IRF7, for instance, were shown to be required for IFN-β induction in response to West Nile virus infection in murine fibroblasts but not in macrophages and dendritic cells, implying that detection of viral components proceeds via different pathways in these cell types [134]. Myeloid cell responses to CMV infection may thus differ from those of fibroblasts, while in latently-infected monocytes, IFN-related gene expression may remain high due to the absence of viral lytic proteins.

Finally, cell responses may also be affected by the virus strain, as virion content of pp65, reported to block IFN induction at very early times post-entry [113, 135], was shown to vary dramatically in different CMV strains [136], while STAT2 degradation was observed to occur in fibroblasts infected with CMV clinical isolates or with strain AD169, but not strain Towne [128]. Altogether, our data broaden the number of IFN-related genes susceptible to transcriptional regulation by CMV to include effectors with currently no known role in CMV infection inhibition, which may thus represent new host encoded anti- or pro-viral proteins.

In summary, our data provide evidence in favor of the existence of a new type of myeloid cells potentially permissive to CMV lytic infection, offer a reasonable theory regarding their preferential infection over other cell types present in the same population, substantially expand our understanding of the cellular determinants of CMV tropism for myeloid and other types of cells, and provide new candidate pro- and anti-viral molecules for future studies and potential therapeutic interventions.

## MATERIALS AND METHODS

### Cells and virus

Umbilical cord blood CD34^+^ HSC were purchased from STEMCELL Technologies Inc, Vancouver, Canada and pre-amplified in α-Minimum Essential Medium (Thermo Fisher Scientific, Waltham, MA) supplemented with 20% heat-inactivated FBS (Gibco, Fisher Scientific, Waltham, MA), 375 ng/ml of FL, 50 ng/ml of SCF and 50 ng/ml of TPO for 8-10 days at a density of 1 × 10^4^ cells/well in 48-well tissue culture plates. Cells were then differentiated in serum-free X-VIVO 15 medium (Lonza/BioWhittaker, Allendale, NJ) supplemented with 1,500 IU/ml of GM-CSF (Leukine Sargramostim), 150 ng/ml of FL, 10 ng/ml of SCF, 2.5 ng/ml of tumor necrosis factor-α, and 0.5 ng/ml of transforming growth factor β1 for eight days at a density of 1 × 10^5^ cells/well in 48-well plates. Activation of differentiated cells was then induced by exposure to X-VIVO 15 medium containing 10% standard FBS (US origin, Gibco, Fisher Scientific, Waltham, MA), 1,500 IU/ml of GM-CSF, 200 ng/ml of CD40L, and 500 ng/ml of LPS (Sigma-Aldrich, St. Louis, MO) for two days at a density of 1 × 10^5^ cells/well in 48-well plates. All cytokines were from Peprotech, Rocky Hill, NJ. Human foreskin fibroblasts were propagated in Dulbecco’s Modified Eagle Medium (Corning Cellgro, UCSF CCF, San Francisco, CA) supplemented with 10% fetal clone serum III, 100 U/ml penicillin, 100 µg/ml streptomycin, 4 mM HEPES (all from HyClone, Fisher Scientific, Pittsburgh, PA), and 1 mM sodium pyruvate (Corning Cellgro, UCSF CCF, San Francisco, CA). CMV strain TB40/E, a gift from C. Sinzger (University of Ulm, Ulm, Germany), was propagated on fibroblasts and purified by ultracentrifugation as previously described [53].

### Myeloid cell infection

Differentiated myeloid cell populations were exposed to TB40/E at a calculated MOI of ten for four hours, washed twice and further cultured for ten days. Cells were harvested on days 2, 4, 6, 8, and 10 pi, counted, and used in immunofluorescence staining analyses and titration assays.

### Immunofluorescence staining analyses

Cell staining was performed as previously described [16]. Briefly, cytospin preparations of myeloid cells were fixed in 1.5% formaldehyde for 30 min, permeabilized in 0.5% Triton-X 100 for 20 min, and blocked in 40% FBS/40% goat serum for 30 min before incubation with antibodies directed against the viral proteins IE1/IE2 (MAb810, 1:600, or AF488 MAB810X, 1:200, Millipore, Temecula, CA), UL32 (pp150, 1:400, a kind gift from Bill Britt, University of Alabama, Birmingham), UL84 (1:500, Virusys, Taneytown, MD), UL44 (1:200, Virusys, Taneytown, MD), or UL57 (1:100, Virusys, Taneytown, MD) for one hour, followed by secondary antibodies conjugated to Alexa-Fluor 488 or Alexa-Fluor 594 (1:200, Invitrogen, Carlsbad, CA, and Jackson Immunoresearch, West Grove, PA) for another hour. Nuclei were labeled with Hoechst 33342 (0.2 mg/ml; Molecular Probes, Eugene, OR) for three min. Samples were viewed using a Nikon Eclipse E600 fluorescence microscope equipped with Ocular imaging software.

### Virus titrations

Cell-associated virus was released from pelleted myeloid cells by sonication for ∼ 5-10 seconds on ice using a Branson Ultrasonics Sonifier 150 and incubated with fibroblasts for one hour. After 24 hours infected fibroblasts were stained for IE1/IE2 expression.

### Statistical analysis

All data were analyzed using Prism 7 (GraphPad Software). Unpaired t-tests were used to compare data from non-activated and activated cells in Fig 3. Differences were considered significant at P < 0.05. The Wilcoxon signed rank sum test was used to compare median ratio values from data distributions with a hypothetical median of zero.

### Single-cell RNA-seq generation and analysis

Activated myeloid cells differentiated from the CD34^+^ HSC of a representative donor (113G) were infected with TB40/E at an MOI of 10, washed twice, and further incubated for 24 hours. Cells were then processed through the Chromium Single-cell 3′ v2 Library Kit (10X Genomics) by the Genetic Resources Core Facility Cell Center and BioRepository, Johns Hopkins University, Baltimore, MD. Briefly, 10,000 cells were loaded onto a single channel of the 10X Chromium Controller. Messenger RNA from approximately ∼ 7,000 cells captured and lysed within nanoliter-sized gel beads in emulsion was then reverse transcribed and barcoded using polyA primers with unique molecular identifier sequences before being pooled, amplified, and used for library preparation. The library was then sequenced in two lanes of an Illumina HiSeq 2500 Rapid Flowcell system. Demultiplexing of the bcl file into a FASTQ file was performed using Cell Ranger mkfastq software, and alignments to human (hg19) or TB40E (NCBI EF999921.1) genome reference sequences were performed using STAR [137]. Dimensionality reduction of data was performed by principal component analysis using N= 10 principal components, and reduced data were visualized in two dimensions using the *t*-SNE nonlinear dimensionality reduction method [18]. Clustering for expression similarity was performed using both graph-based and K-means (with K=10 clusters) methods by Cell Ranger [42]. Clusters and differential expression analyses generated by Cell Ranger were then visualized using Loupe^TM^ Cell Browser [19]. For each gene in each cluster, three values were computed and reported in supplemental datasets: 1) the mean number of unique molecular identifier counts; 2) the log2 fold-change of each gene’s expression in cluster × relative to other clusters and 3) the p-value denoting significance of each gene’s expression in cluster × relative to other clusters, adjusted to account for the number of hypotheses (*i.e.*, genes) being tested.

### Monocle clustering and single cell ordering in pseudotime

Cells belonging to the cluster 7, erythro, mono, MDDC, CMV^+^, promyelo, act neut, and sub-cluster 3 groups depicted in Fig 4C were used for pseudotime analysis. Gene-cell matrices produced by Cell Ranger were loaded into R with cellrangerRkit (https://support.10xgenomics.com/single-cell-gene-expression/software/pipelines/latest/rkit) and pseudo-temporal assignment was performed with Monocle version 2.99.0 (39) using N = 5 principal components. Marker genes were found using Seurat’s FindAllMarkers function [52], and groups were identified based on the expression of gene markers from Fig 4C and S6 Fig. The root of the tree was manually selected using orderCells from Monocle, defined by the point of origin of the majority of the branches.

### Data availability

All single-cell data files are deposited in Gene Expression Omnibus under accession number GSExx.

## SUPPORTING INFORMATION

**S1 Dataset.** Cellular genes differentially expressed in CMV-transcript^+^ versus CMV-transcript-cells at P < 0.05.

**S2 Dataset.** Full name and symbol of genes mentioned in the text.

**S3 Dataset.** Cellular genes with four-fold higher mean expression levels in CMV^−^ than in CMV^+^ cells and present in more than 50% of CMV^−^ cells, but less than 50% of CMV^+^ cells.

**S4 Fig.** Transcript abundance and distribution of the nine cellular genes with four-fold higher mean expression levels in CMV^−^ than in CMV^+^ cells and present in more than 50% of CMV^−^ cells, but less than 50% of CMV^+^ cells.

*t*-SNE projection of data from profiled cells colored based on their quantitative (Log_2_ Gene Exp Max) content in transcripts mapping to the cellular genes named in each box. The proportion of CMV^+^ and CMV^−^ cells expressing each gene is indicated beside the CMV^+^ cluster and in the bottom right corner of each box, respectively.

**S5 Dataset.** Cellular genes significantly enriched (Log_2_ fold change > 4, P values < 10^−15^) in cluster 6 relative to the rest of the population.

**S6 Fig.** Transcript abundance and distribution of promyelocytes, activated neutrophils, and sub-cluster 3 cellular gene markers. *t*-SNE projection of data from profiled cells colored based on their quantitative (Log_2_ Gene Exp Max) content in transcripts mapping to the genes listed in the lower left corner of each panel and corresponding to the promyelocytes (A), activated neutrophils (B), or sub-cluster 3/GEMM (C) clusters in Fig 4C.

**S7 Dataset.** Expression distribution of the 115 cellular genes characterizing sub-cluster 3 as compared to data from Velten L. et al and Karamitros D. et al.

**S8 Fig.** Transcript abundance of gene markers used to characterize the cell groups shown in Fig 5. Monocle pseudotime trajectory of cells colored based on their content in transcripts mapping to the genes listed in the upper left corner of each panel and corresponding to the GR A – MDDC (A), GR B – CL7 (B), GR C – CMV (C), GR D – GEMM (D), GR E – Mono (E), GR F – Erythro (F), GR G – Act Neut (G), and GR H – Promyelo (H) groups in Fig 5B.

**S9 Dataset.** Cellular genes differentially expressed in each of the eight groups generated by Monocle as depicted in Fig 5.

**S10 Dataset.** Cellular genes differentially expressed in GEMM and CMV^+^ cells relative to the rest of the population and their partitioning into functional categories.

## ACKNOWLEDGMENTS

We thank Melissa Olson, Director, Genetics Research Core Facility and Biorepository & Cell Center and David Mohr, Director, High Throughput Sequencing at the Johns Hopkins Medical Institutes, Baltimore MD for technical assistance with single-cell RNA-seq analyses using the 10X genomics Chromium platform. We are also grateful to Aharon Nachshon, Weizmann Institute of Science, for sharing the annotated reference for the TB40/E transcription units, to Christian Sinzger for the kind gift of CMV strain TB40/E, and to Bill Britt for providing us with the anti-pp150 antibodies.

## REFERENCES

1. Griffiths P, Plotkin S, Mocarski E, Pass R, Schleiss M, Krause P, et al. Desirability and feasibility of a vaccine against cytomegalovirus. Vaccine. 2013;31 Suppl 2:B197–203. Epub 2013/04/26. doi: S0264-410X(12)01534-4 [pii] 10.1016/j.vaccine.2012.10.074. PubMed PMID: 23598482.

2. Pass RF. Cytomegalovirus. In: Howley DKaP, editor. Fields virology. Philadelphia: Lippincott/The Williams & Wilkins Co.; 2001. p. 2675–705.

3. Dupont L, Reeves MB. Cytomegalovirus latency and reactivation: recent insights into an age old problem. Rev Med Virol. 2016;26(2):75–89. doi: 10.1002/rmv.1862. PubMed PMID: 26572645; PubMed Central PMCID: PMCPMC5458136.

4. Sinclair J, Reeves M. The intimate relationship between human cytomegalovirus and the dendritic cell lineage. Frontiers in microbiology. 2014;5:389. doi: 10.3389/fmicb.2014.00389. PubMed PMID: 25147545; PubMed Central PMCID: PMCPMC4124589.

5. Hertel L. Human cytomegalovirus tropism for mucosal myeloid dendritic cells. Rev Med Virol. 2014;24(6):379–95. doi: 10.1002/rmv.1797. PubMed PMID: 24888709; PubMed Central PMCID: PMCPMC4213286.

6. Stevenson EV, Collins-McMillen D, Kim JH, Cieply SJ, Bentz GL, Yurochko AD. HCMV reprogramming of infected monocyte survival and differentiation: a Goldilocks phenomenon. Viruses. 2014;6(2):782–807. doi: 10.3390/v6020782. PubMed PMID: 24531335; PubMed Central PMCID: PMCPMC3939482.

7. Kondo K, Kaneshima H, Mocarski ES. Human cytomegalovirus latent infection of granulocyte-macrophage progenitors. Proc Natl Acad Sci U S A. 1994;91(25):11879–83. PubMed PMID: 7991550.

8. Goodrum FD, Jordan CT, High K, Shenk T. Human cytomegalovirus gene expression during infection of primary hematopoietic progenitor cells: a model for latency. Proc Natl Acad Sci U S A. 2002;99(25):16255–60. PubMed PMID: 12456880.

9. Hertel L, Lacaille VG, Strobl H, Mellins ED, Mocarski ES. Susceptibility of immature and mature Langerhans cell-type dendritic cells to infection and immunomodulation by human cytomegalovirus. J Virol. 2003;77(13):7563–74. Epub 2003/06/14. PubMed PMID: 12805456; PubMed Central PMCID: PMC164783.

10. Reeves MB, Lehner PJ, Sissons JG, Sinclair JH. An in vitro model for the regulation of human cytomegalovirus latency and reactivation in dendritic cells by chromatin remodelling. J Gen Virol. 2005;86(Pt 11):2949–54. PubMed PMID: 16227215.

11. Ibanez CE, Schrier R, Ghazal P, Wiley C, Nelson JA. Human cytomegalovirus productively infects primary differentiated macrophages. J Virol. 1991;65(12):6581–8. PubMed PMID: 1658363.

12. Strobl H, Bello-Fernandez C, Riedl E, Pickl WF, Majdic O, Lyman SD, et al. flt3 ligand in cooperation with transforming growth factor-beta1 potentiates in vitro development of Langerhans-type dendritic cells and allows single-cell dendritic cell cluster formation under serum-free conditions. Blood. 1997;90(4):1425–34. PubMed PMID: 9269760.

13. Strobl H, Krump C, Borek I. Micro-environmental signals directing human epidermal Langerhans cell differentiation. Seminars in cell & developmental biology. 2018. doi: 10.1016/j.semcdb.2018.02.016. PubMed PMID: 29448069.

14. Lauron EJ, Yu D, Fehr AR, ertel L. Human cytomegalovirus infection of langerhans-type dendritic cells does not require the presence of the gH/gL/UL128-131A complex and is blocked after nuclear deposition of viral genomes in immature cells. J Virol. 2014;88(1):403–16. Epub 2013/10/25. doi: 10.1128/JVI.03062-13. PubMed PMID: 24155395; PubMed Central PMCID: PMC3911714.

15. Coronel R, Takayama S, Juwono T, Hertel L. Dynamics of Human Cytomegalovirus Infection in CD34+ Hematopoietic Cells and Derived Langerhans-Type Dendritic Cells. J Virol. 2015;89(10):5615–32. Epub 2015/03/13. doi: 10.1128/JVI.00305-15. PubMed PMID: 25762731; PubMed Central PMCID: PMC4442541.

16. Coronel R, Jesus DM, Dalle Ore L, Mymryk JS, Hertel L. Activation of Langerhans-Type Dendritic Cells Alters Human Cytomegalovirus Infection and Reactivation in a Stimulus-Dependent Manner. Frontiers in microbiology. 2016;7:1445. doi: 10.3389/fmicb.2016.01445. PubMed PMID: 27683575; PubMed Central PMCID: PMCPMC5021960.

17. 10XGenomicsChromium. https://www.10xgenomics.com/solutions/single-cell/. 2018.

18. van der Maaten LJP, Hinton GE. Visualizing High-Dimensional Data Using t-SNE. Journal of Machine Learning Research. 2008;9:2579–605.

19. CellBrowser. https://support.10xgenomics.com/single-cell-gene-expression/software/visualization/latest/what-is-loupe-cell-browser. 2018.

20. Goodrum F, Jordan CT, Terhune SS, High K, Shenk T. Differential outcomes of human cytomegalovirus infection in primitive hematopoietic cell subpopulations. Blood. 2004;104(3):687–95. PubMed PMID: 15090458.

21. Bresnahan WA, Shenk TE. UL82 virion protein activates expression of immediate early viral genes in human cytomegalovirus-infected cells. Proc Natl Acad Sci U S A. 2000;97(26):14506–11. PubMed PMID: 11121054.

22. Greijer AE, Dekkers CA, Middeldorp JM. Human cytomegalovirus virions differentially incorporate viral and host cell RNA during the assembly process. J Virol. 2000;74(19):9078–82. PubMed PMID: 10982353; PubMed Central PMCID: PMCPMC102105.

23. Ogawa-Goto K, Tanaka K, Gibson W, Moriishi E, Miura Y, Kurata T, et al. Microtubule network facilitates nuclear targeting of human cytomegalovirus capsid. J Virol. 2003;77(15):8541–7. PubMed PMID: 12857923.

24. Miller MS, Hertel L. Onset of human cytomegalovirus replication in fibroblasts requires the presence of an intact vimentin cytoskeleton. J Virol. 2009;83(14):7015–28. Epub 2009/05/01. doi: JVI.00398-09 [pii] 10.1128/JVI.00398-09. PubMed PMID: 19403668; PubMed Central PMCID: PMC2704777.

25. Wang X, Huong SM, Chiu ML, Raab-Traub N, Huang ES. Epidermal growth factor receptor is a cellular receptor for human cytomegalovirus. Nature. 2003;424(6947):456–61. PubMed PMID: 12879076.

26. Chan G, Nogalski MT, Yurochko AD. Activation of EGFR on monocytes is required for human cytomegalovirus entry and mediates cellular motility. Proc Natl Acad Sci U S A. 2009;106(52):22369–74. Epub 2009/12/19. doi: 0908787106 [pii] 10.1073/pnas.0908787106. PubMed PMID: 20018733; PubMed Central PMCID: PMC2799688.

27. Kim JH, Collins-McMillen D, Buehler JC, Goodrum FD, Yurochko AD. Human Cytomegalovirus Requires Epidermal Growth Factor Receptor Signaling To Enter and Initiate the Early Steps in the Establishment of Latency in CD34(+) Human Progenitor Cells. J Virol. 2017;91(5). doi: 10.1128/JVI.01206-16. PubMed PMID: 27974567; PubMed Central PMCID: PMCPMC5309964.

28. Buehler J, Zeltzer S, Reitsma J, Petrucelli A, Umashankar M, Rak M, et al. Opposing Regulation of the EGF Receptor: A Molecular Switch Controlling Cytomegalovirus Latency and Replication. PLoS Pathog. 2016;12(5):e1005655. doi: 10.1371/journal.ppat.1005655. PubMed PMID: 27218650; PubMed Central PMCID: PMCPMC4878804.

29. Soroceanu L, Akhavan A, Cobbs CS. Platelet-derived growth factor-alpha receptor activation is required for human cytomegalovirus infection. Nature. 2008;455(7211):391–5. PubMed PMID: 18701889.

30. Kabanova A, Marcandalli J, Zhou T, Bianchi S, Baxa U, Tsybovsky Y, et al. Platelet-derived growth factor-alpha receptor is the cellular receptor for human cytomegalovirus gHgLgO trimer. Nat Microbiol. 2016;1(8):16082. doi: 10.1038/nmicrobiol.2016.82. PubMed PMID: 27573107; PubMed Central PMCID: PMCPMC4918640.

31. Wu Y, Prager A, Boos S, Resch M, Brizic I, Mach M, et al. Human cytomegalovirus glycoprotein complex gH/gL/gO uses PDGFR-alpha as a key for entry. PLoS Pathog. 2017;13(4):e1006281. doi: 10.1371/journal.ppat.1006281. PubMed PMID: 28403202; PubMed Central PMCID: PMCPMC5389851.

32. Li Q, Wilkie AR, Weller M, Liu X, Cohen JI. THY-1 Cell Surface Antigen (CD90) Has an Important Role in the Initial Stage of Human Cytomegalovirus Infection. PLoS Pathog. 2015;11(7):e1004999. doi: 10.1371/journal.ppat.1004999. PubMed PMID: 26147640; PubMed Central PMCID: PMCPMC4492587.

33. Li Q, Fischer E, Cohen JI. Cell Surface THY-1 Contributes to Human Cytomegalovirus Entry via a Macropinocytosis-Like Process. J Virol. 2016;90(21):9766–81. doi: 10.1128/JVI.01092-16. PubMed PMID: 27558416; PubMed Central PMCID: PMCPMC5068528.

34. Feire AL, Koss H, Compton T. Cellular integrins function as entry receptors for human cytomegalovirus via a highly conserved disintegrin-like domain. Proc Natl Acad Sci U S A. 2004;101(43):15470–5. PubMed PMID: 15494436.

35. Feire AL, Roy RM, Manley K, Compton T. The glycoprotein B disintegrin-like domain binds beta 1 integrin to mediate cytomegalovirus entry. J Virol. 2010;84(19):10026–37. doi: 10.1128/JVI.00710-10. PubMed PMID: 20660204; PubMed Central PMCID: PMCPMC2937812.

36. Wang X, Huang DY, Huong SM, Huang ES. Integrin alphavbeta3 is a coreceptor for human cytomegalovirus. Nat Med. 2005;11(5):515–21. PubMed PMID: 15834425.

37. Vanarsdall AL, Pritchard SR, Wisner TW, Liu J, Jardetzky TS, Johnson DC. CD147 Promotes Entry of Pentamer-Expressing Human Cytomegalovirus into Epithelial and Endothelial Cells. MBio. 2018;9(3). doi: 10.1128/mBio.00781-18. PubMed PMID: 29739904; PubMed Central PMCID: PMCPMC5941078.

38. Pari GS, Anders DG. Eleven loci encoding trans-acting factors are required for transient complementation of human cytomegalovirus oriLyt-dependent DNA replication. J Virol. 1993;67(12):6979–88. PubMed PMID: 8230421; PubMed Central PMCID: PMCPMC238157.

39. Pari GS. Nuts and bolts of human cytomegalovirus lytic DNA replication. Curr Top Microbiol Immunol. 2008;325:153–66. PubMed PMID: 18637505.

40. Boehm T, Hofer S, Winklehner P, Kellersch B, Geiger C, Trockenbacher A, et al. Attenuation of cell adhesion in lymphocytes is regulated by CYTIP, a protein which mediates signal complex sequestration. EMBO J. 2003;22(5):1014–24. doi: 10.1093/emboj/cdg101. PubMed PMID: 12606567; PubMed Central PMCID: PMCPMC150334.

41. Ranheim EA, Kipps TJ. Activated T cells induce expression of B7/BB1 on normal or leukemic B cells through a CD40-dependent signal. J Exp Med. 1993;177(4):925–35. PubMed PMID: 7681471; PubMed Central PMCID: PMCPMC2190967.

42. CellRanger. https://support.10xgenomics.com/single-cell-gene-expression/software/pipelines/latest/what-is-cell-ranger. 2018.

43. Rapin N, Bagger FO, Jendholm J, Mora-Jensen H, Krogh A, Kohlmann A, et al. Comparing cancer vs normal gene expression profiles identifies new disease entities and common transcriptional programs in AML patients. Blood. 2014;123(6):894–904. doi: 10.1182/blood-2013-02-485771. PubMed PMID: 24363398.

44. Mabbott NA, Baillie JK, Brown H, Freeman TC, Hume DA. An expression atlas of human primary cells: inference of gene function from coexpression networks. BMC Genomics. 2013;14:632. doi: 10.1186/1471-2164-14-632. PubMed PMID: 24053356; PubMed Central PMCID: PMCPMC3849585.

45. Tardif MR, Chapeton-Montes JA, Posvandzic A, Page N, Gilbert C, Tessier PA. Secretion of S100A8, S100A9, and S100A12 by Neutrophils Involves Reactive Oxygen Species and Potassium Efflux. J Immunol Res. 2015;2015:296149. doi: 10.1155/2015/296149. PubMed PMID: 27057553; PubMed Central PMCID: PMCPMC4736198.

46. Bostrom EA, Tarkowski A, Bokarewa M. Resistin is stored in neutrophil granules being released upon challenge with inflammatory stimuli. Biochim Biophys Acta. 2009;1793(12):1894–900. doi: 10.1016/j.bbamcr.2009.09.008. PubMed PMID: 19770005.

47. Karamitros D, Stoilova B, Aboukhalil Z, Hamey F, Reinisch A, Samitsch M, et al. Single-cell analysis reveals the continuum of human lympho-myeloid progenitor cells. Nat Immunol. 2018;19(1):85–97. doi: 10.1038/s41590-017-0001-2. PubMed PMID: 29167569; PubMed Central PMCID: PMCPMC5884424.

48. Velten L, Haas SF, Raffel S, Blaszkiewicz S, Islam S, Hennig BP, et al. Human haematopoietic stem cell lineage commitment is a continuous process. Nat Cell Biol. 2017;19(4):271–81. doi: 10.1038/ncb3493. PubMed PMID: 28319093; PubMed Central PMCID: PMCPMC5496982.

49. Laurenti E, Gottgens B. From haematopoietic stem cells to complex differentiation landscapes. Nature. 2018;553(7689):418–26. doi: 10.1038/nature25022. PubMed PMID: 29364285.

50. Cabezas-Wallscheid N, Klimmeck D, Hansson J, Lipka DB, Reyes A, Wang Q, et al. Identification of regulatory networks in HSCs and their immediate progeny via integrated proteome, transcriptome, and DNA methylome analysis. Cell Stem Cell. 2014;15(4):507–22. doi: 10.1016/j.stem.2014.07.005. PubMed PMID: 25158935.

51. Trapnell C, Cacchiarelli D, Grimsby J, Pokharel P, Li S, Morse M, et al. The dynamics and regulators of cell fate decisions are revealed by pseudotemporal ordering of single cells. Nat Biotechnol. 2014;32(4):381–6. doi: 10.1038/nbt.2859. PubMed PMID: 24658644; PubMed Central PMCID: PMCPMC4122333.

52. Butler A, Hoffman P, Smibert P, Papalexi E, Satija R. Integrating single-cell transcriptomic data across different conditions, technologies, and species. Nat Biotechnol. 2018;36(5):411–20. doi: 10.1038/nbt.4096. PubMed PMID: 29608179.

53. Hertel L, Mocarski ES. Global analysis of host cell gene expression late during cytomegalovirus infection reveals extensive dysregulation of cell cycle gene expression and induction of Pseudomitosis independent of US28 function. J Virol. 2004;78(21):11988–2011. PubMed PMID: 15479839.

54. Bresnahan WA, Boldogh I, Thompson EA, Albrecht T. Human cytomegalovirus inhibits cellular DNA synthesis and arrests productively infected cells in late G1. Virology. 1996;224(1):150–60. PubMed PMID: 8862409.

55. Dittmer D, Mocarski ES. Human cytomegalovirus infection inhibits G1/S transition. J Virol. 1997;71(2):1629–34. PubMed PMID: 8995690.

56. Lu M, Shenk T. Human cytomegalovirus infection inhibits cell cycle progression at multiple points, including the transition from G1 to S. J Virol. 1996;70(12):8850–7. PubMed PMID: 8971013.

57. Asai T, Liu Y, Di Giandomenico S, Bae N, Ndiaye-Lobry D, Deblasio A, et al. Necdin, a p53 target gene, regulates the quiescence and response to genotoxic stress of hematopoietic stem/progenitor cells. Blood. 2012;120(8):1601–12. doi: 10.1182/blood-2011-11-393983. PubMed PMID: 22776820; PubMed Central PMCID: PMCPMC3429304.

58. Hertel L, Chou S, Mocarski ES. Viral and cell cycle-regulated kinases in cytomegalovirus-induced pseudomitosis and replication. PLoS Pathog. 2007;3(1):e6. PubMed PMID: 17206862.

59. Kabir MA, Uddin W, Narayanan A, Reddy PK, Jairajpuri MA, Sherman F, et al. Functional Subunits of Eukaryotic Chaperonin CCT/TRiC in Protein Folding. J Amino Acids. 2011;2011:843206. doi: 10.4061/2011/843206. PubMed PMID: 22312474; PubMed Central PMCID: PMCPMC3268035.

60. Meunier L, Usherwood YK, Chung KT, Hendershot LM. A subset of chaperones and folding enzymes form multiprotein complexes in endoplasmic reticulum to bind nascent proteins. Mol Biol Cell. 2002;13(12):4456–69. doi: 10.1091/mbc.e02-05-0311. PubMed PMID: 12475965; PubMed Central PMCID: PMCPMC138646.

61. Williams DB. Beyond lectins: the calnexin/calreticulin chaperone system of the endoplasmic reticulum. J Cell Sci. 2006;119(Pt 4):615–23. doi: 10.1242/jcs.02856. PubMed PMID: 16467570.

62. Khan S, Zimmermann A, Basler M, Groettrup M, Hengel H. A cytomegalovirus inhibitor of gamma interferon signaling controls immunoproteasome induction. J Virol. 2004;78(4):1831–42. PubMed PMID: 14747547; PubMed Central PMCID: PMCPMC369451.

63. Tuli A, Sharma M, McIlhaney MM, Talmadge JE, Naslavsky N, Caplan S, et al. Amyloid precursor-like protein 2 increases the endocytosis, instability, and turnover of the H2-K(d) MHC class I molecule. J Immunol. 2008;181(3):1978–87. PubMed PMID: 18641335; PubMed Central PMCID: PMCPMC2607064.

64. Tuli A, Sharma M, Naslavsky N, Caplan S, Solheim JC. Specificity of amyloid precursor-like protein 2 interactions with MHC class I molecules. Immunogenetics. 2008;60(6):303–13. doi: 10.1007/s00251-008-0296-0. PubMed PMID: 18452037; PubMed Central PMCID: PMCPMC2683759.

65. ten Broeke T, Wubbolts R, Stoorvogel W. MHC class II antigen presentation by dendritic cells regulated through endosomal sorting. Cold Spring Harb Perspect Biol. 2013;5(12):a016873. doi: 10.1101/cshperspect.a016873. PubMed PMID: 24296169; PubMed Central PMCID: PMCPMC3839614.

66. Unterholzner L, Keating SE, Baran M, Horan KA, Jensen SB, Sharma S, et al. IFI16 is an innate immune sensor for intracellular DNA. Nat Immunol. 2010;11(11):997–1004. doi: 10.1038/ni.1932. PubMed PMID: 20890285; PubMed Central PMCID: PMCPMC3142795.

67. Yanai H, Ban T, Wang Z, Choi MK, Kawamura T, Negishi H, et al. HMGB proteins function as universal sentinels for nucleic-acid-mediated innate immune responses. Nature. 2009;462(7269):99–103. doi: 10.1038/nature08512. PubMed PMID: 19890330.

68. Pichlmair A, Schulz O, Tan CP, Naslund TI, Liljestrom P, Weber F, et al. RIG-I-mediated antiviral responses to single-stranded RNA bearing 5’-phosphates. Science. 2006;314(5801):997–1001. doi: 10.1126/science.1132998. PubMed PMID: 17038589.

69. Kato H, Takeuchi O, Sato S, Yoneyama M, Yamamoto M, Matsui K, et al. Differential roles of MDA5 and RIG-I helicases in the recognition of RNA viruses. Nature. 2006;441(7089):101–5. doi: 10.1038/nature04734. PubMed PMID: 16625202.

70. Meurs E, Chong K, Galabru J, Thomas NS, Kerr IM, Williams BR, et al. Molecular cloning and characterization of the human double-stranded RNA-activated protein kinase induced by interferon. Cell. 1990;62(2):379–90. PubMed PMID: 1695551.

71. Zhao GN, Jiang DS, Li H. Interferon regulatory factors: at the crossroads of immunity, metabolism, and disease. Biochim Biophys Acta. 2015;1852(2):365–78. doi: 10.1016/j.bbadis.2014.04.030. PubMed PMID: 24807060.

72. Negishi H, Ohba Y, Yanai H, Takaoka A, Honma K, Yui K, et al. Negative regulation of Toll-like-receptor signaling by IRF-4. Proc Natl Acad Sci U S A. 2005;102(44):15989–94. doi: 10.1073/pnas.0508327102. PubMed PMID: 16236719; PubMed Central PMCID: PMCPMC1257749.

73. Harada H, Fujita T, Miyamoto M, Kimura Y, Maruyama M, Furia A, et al. Structurally similar but functionally distinct factors, IRF-1 and IRF-2, bind to the same regulatory elements of IFN and IFN-inducible genes. Cell. 1989;58(4):729–39. PubMed PMID: 2475256.

74. Sanada T, Takaesu G, Mashima R, Yoshida R, Kobayashi T, Yoshimura A. FLN29 deficiency reveals its negative regulatory role in the Toll-like receptor (TLR) and retinoic acid-inducible gene I (RIG-I)-like helicase signaling pathway. J Biol Chem. 2008;283(49):33858–64. doi: 10.1074/jbc.M806923200. PubMed PMID: 18849341; PubMed Central PMCID: PMCPMC2662213.

75. Yoshimura A, Ito M, Chikuma S, Akanuma T, Nakatsukasa H. Negative Regulation of Cytokine Signaling in Immunity. Cold Spring Harb Perspect Biol. 2018;10(7). doi: 10.1101/cshperspect.a028571. PubMed PMID: 28716890.

76. Vladimer GI, Gorna MW, Superti-Furga G. IFITs: Emerging Roles as Key Anti-Viral Proteins. Front Immunol. 2014;5:94. doi: 10.3389/fimmu.2014.00094. PubMed PMID: 24653722; PubMed Central PMCID: PMCPMC3948006.

77. Shi G, Schwartz O, Compton AA. More than meets the I: the diverse antiviral and cellular functions of interferon-induced transmembrane proteins. Retrovirology. 2017;14(1):53. doi: 10.1186/s12977-017-0377-y. PubMed PMID: 29162141; PubMed Central PMCID: PMCPMC5697417.

78. Perng YC, Lenschow DJ. ISG15 in antiviral immunity and beyond. Nat Rev Microbiol. 2018;16(7):423–39. doi: 10.1038/s41579-018-0020-5. PubMed PMID: 29769653.

79. Haller O, Staeheli P, Schwemmle M, Kochs G. Mx GTPases: dynamin-like antiviral machines of innate immunity. Trends Microbiol. 2015;23(3):154–63. doi: 10.1016/j.tim.2014.12.003. PubMed PMID: 25572883.

80. Hovanessian AG, Justesen J. The human 2’-5’oligoadenylate synthetase family: unique interferon-inducible enzymes catalyzing 2’-5’ instead of 3’-5’ phosphodiester bond formation. Biochimie. 2007;89(6-7):779–88. doi: 10.1016/j.biochi.2007.02.003. PubMed PMID: 17408844.

81. Helbig KJ, Beard MR. The role of viperin in the innate antiviral response. J Mol Biol. 2014;426(6):1210–9. doi: 10.1016/j.jmb.2013.10.019. PubMed PMID: 24157441.

82. Li M, Zhang D, Zhu M, Shen Y, Wei W, Ying S, et al. Roles of SAMHD1 in antiviral defense, autoimmunity and cancer. Rev Med Virol. 2017;27(4). doi: 10.1002/rmv.1931. PubMed PMID: 28444859.

83. Zheng Z, Wang L, Pan J. Interferon-stimulated gene 20-kDa protein (ISG20) in infection and disease: Review and outlook. Intractable Rare Dis Res. 2017;6(1):35–40. doi: 10.5582/irdr.2017.01004. PubMed PMID: 28357179; PubMed Central PMCID: PMCPMC5359350.

84. Ishikawa H, Barber GN. STING is an endoplasmic reticulum adaptor that facilitates innate immune signalling. Nature. 2008;455(7213):674–8. doi: 10.1038/nature07317. PubMed PMID: 18724357; PubMed Central PMCID: PMCPMC2804933.

85. Ishikawa H, Ma Z, Barber GN. STING regulates intracellular DNA-mediated, type I interferon-dependent innate immunity. Nature. 2009;461(7265):788–92. doi: 10.1038/nature08476. PubMed PMID: 19776740; PubMed Central PMCID: PMCPMC4664154.

86. Tanaka Y, Chen ZJ. STING specifies IRF3 phosphorylation by TBK1 in the cytosolic DNA signaling pathway. Sci Signal. 2012;5(214):ra20. doi: 10.1126/scisignal.2002521. PubMed PMID: 22394562; PubMed Central PMCID: PMCPMC3549669.

87. Zhong B, Yang Y, Li S, Wang YY, Li Y, Diao F, et al. The adaptor protein MITA links virus-sensing receptors to IRF3 transcription factor activation. Immunity. 2008;29(4):538–50. doi: 10.1016/j.immuni.2008.09.003. PubMed PMID: 18818105.

88. Outlioua A, Pourcelot M, Arnoult D. The Role of Optineurin in Antiviral Type I Interferon Production. Front Immunol. 2018;9:853. doi: 10.3389/fimmu.2018.00853. PubMed PMID: 29755463; PubMed Central PMCID: PMCPMC5932347.

89. Paijo J, Doring M, Spanier J, Grabski E, Nooruzzaman M, Schmidt T, et al. cGAS Senses Human Cytomegalovirus and Induces Type I Interferon Responses in Human Monocyte-Derived Cells. PLoS Pathog. 2016;12(4):e1005546. doi: 10.1371/journal.ppat.1005546. PubMed PMID: 27058035; PubMed Central PMCID: PMCPMC4825940.

90. Shnayder M, Nachshon A, Krishna B, Poole E, Boshkov A, Binyamin A, et al. Defining the Transcriptional Landscape during Cytomegalovirus Latency with Single-Cell RNA Sequencing. MBio. 2018;9(2). doi: 10.1128/mBio.00013-18. PubMed PMID: 29535194; PubMed Central PMCID: PMCPMC5850328s.

91. Waddington CH. The strategy of the genes: a discussion of some aspects of theoretical biology: Allen & Unwin; 1957.

92. Povinelli BJ, Rodriguez-Meira A, Mead AJ. Single cell analysis of normal and leukemic hematopoiesis. Mol Aspects Med. 2018;59:85–94. doi: 10.1016/j.mam.2017.08.006. PubMed PMID: 28863981; PubMed Central PMCID: PMCPMC5771467.

93. Buenrostro JD, Corces MR, Lareau CA, Wu B, Schep AN, Aryee MJ, et al. Integrated Single-Cell Analysis Maps the Continuous Regulatory Landscape of Human Hematopoietic Differentiation. Cell. 2018;173(6):1535–48 e16. doi: 10.1016/j.cell.2018.03.074. PubMed PMID: 29706549; PubMed Central PMCID: PMCPMC5989727.

94. Piacibello W, Sanavio F, Garetto L, Severino A, Bergandi D, Ferrario J, et al. Extensive amplification and self-renewal of human primitive hematopoietic stem cells from cord blood. Blood. 1997;89(8):2644–53. Epub 1997/04/15. PubMed PMID: 9108381.

95. Tanavde VM, Malehorn MT, Lumkul R, Gao Z, Wingard J, Garrett ES, et al. Human stem-progenitor cells from neonatal cord blood have greater hematopoietic expansion capacity than those from mobilized adult blood. Exp Hematol. 2002;30(7):816–23. PubMed PMID: 12135681.

96. Gilmore GL, DePasquale DK, Lister J, Shadduck RK. Ex vivo expansion of human umbilical cord blood and peripheral blood CD34(+) hematopoietic stem cells. Exp Hematol. 2000;28(11):1297–305. PubMed PMID: 11063878.

97. Psatha N, Karponi G, Yannaki E. Optimizing autologous cell grafts to improve stem cell gene therapy. Exp Hematol. 2016;44(7):528–39. doi: 10.1016/j.exphem.2016.04.007. PubMed PMID: 27106799; PubMed Central PMCID: PMCPMC4914411.

98. Flores-Guzman P, Fernandez-Sanchez V, Mayani H. Concise review: ex vivo expansion of cord blood-derived hematopoietic stem and progenitor cells: basic principles, experimental approaches, and impact in regenerative medicine. Stem Cells Transl Med. 2013;2(11):830–8. doi: 10.5966/sctm.2013-0071. PubMed PMID: 24101670; PubMed Central PMCID: PMCPMC3808198.

99. Machlus KR, Italiano JE, Jr. The incredible journey: From megakaryocyte development to platelet formation. J Cell Biol. 2013;201(6):785–96. doi: 10.1083/jcb.201304054. PubMed PMID: 23751492; PubMed Central PMCID: PMCPMC3678154.

100. Tsapogas P, Mooney CJ, Brown G, Rolink A. The Cytokine Flt3-Ligand in Normal and Malignant Hematopoiesis. Int J Mol Sci. 2017;18(6). doi: 10.3390/ijms18061115. PubMed PMID: 28538663; PubMed Central PMCID: PMCPMC5485939.

101. Munger J, Bajad SU, Coller HA, Shenk T, Rabinowitz JD. Dynamics of the cellular metabolome during human cytomegalovirus infection. PLoS Pathog. 2006;2(12):e132. doi: 10.1371/journal.ppat.0020132. PubMed PMID: 17173481; PubMed Central PMCID: PMCPMC1698944.

102. Kaarbo M, Ager-Wick E, Osenbroch PO, Kilander A, Skinnes R, Muller F, et al. Human cytomegalovirus infection increases mitochondrial biogenesis. Mitochondrion. 2011;11(6):935–45. doi: 10.1016/j.mito.2011.08.008. PubMed PMID: 21907833.

103. Jean Beltran PM, Mathias RA, Cristea IM. A Portrait of the Human Organelle Proteome In Space and Time during Cytomegalovirus Infection. Cell Syst. 2016;3(4):361–73 e6. doi: 10.1016/j.cels.2016.08.012. PubMed PMID: 27641956; PubMed Central PMCID: PMCPMC5083158.

104. Karniely S, Weekes MP, Antrobus R, Rorbach J, van Haute L, Umrania Y, et al. Human Cytomegalovirus Infection Upregulates the Mitochondrial Transcription and Translation Machineries. MBio. 2016;7(2):e00029. doi: 10.1128/mBio.00029-16. PubMed PMID: 27025248; PubMed Central PMCID: PMCPMC4807356.

105. Spector DH. Human cytomegalovirus riding the cell cycle. Med Microbiol Immunol. 2015;204(3):409–19. doi: 10.1007/s00430-015-0396-z. PubMed PMID: 25776080.

106. Eifler M, Uecker R, Weisbach H, Bogdanow B, Richter E, Konig L, et al. PUL21a-Cyclin A2 interaction is required to protect human cytomegalovirus-infected cells from the deleterious consequences of mitotic entry. PLoS Pathog. 2014;10(10):e1004514. doi: 10.1371/journal.ppat.1004514. PubMed PMID: 25393019; PubMed Central PMCID: PMCPMC4231158.

107. Tirosh O, Cohen Y, Shitrit A, Shani O, Le-Trilling VT, Trilling M, et al. The Transcription and Translation Landscapes during Human Cytomegalovirus Infection Reveal Novel Host-Pathogen Interactions. PLoS Pathog. 2015;11(11):e1005288. doi: 10.1371/journal.ppat.1005288. PubMed PMID: 26599541; PubMed Central PMCID: PMCPMC4658056.

108. Palpant NJ, Wang Y, Hadland B, Zaunbrecher RJ, Redd M, Jones D, et al. Chromatin and Transcriptional Analysis of Mesoderm Progenitor Cells Identifies HOPX as a Regulator of Primitive Hematopoiesis. Cell Rep. 2017;20(7):1597–608. doi: 10.1016/j.celrep.2017.07.067. PubMed PMID: 28813672; PubMed Central PMCID: PMCPMC5576510.

109. Ling F, Kang B, Sun XH. Id proteins: small molecules, mighty regulators. Curr Top Dev Biol. 2014;110:189–216. doi: 10.1016/B978-0-12-405943-6.00005-1. PubMed PMID: 25248477.

110. Lotem J, Levanon D, Negreanu V, Bauer O, Hantisteanu S, Dicken J, et al. Runx3 in Immunity, Inflammation and Cancer. Adv Exp Med Biol. 2017;962:369–93. doi: 10.1007/978-981-10-3233-2_23. PubMed PMID: 28299669.

111. Marke R, van Leeuwen FN, Scheijen B. The many faces of IKZF1 in B-cell precursor acute lymphoblastic leukemia. Haematologica. 2018;103(4):565–74. doi: 10.3324/haematol.2017.185603. PubMed PMID: 29519871; PubMed Central PMCID: PMCPMC5865415.

112. Burda P, Laslo P, Stopka T. The role of PU.1 and GATA-1 transcription factors during normal and leukemogenic hematopoiesis. Leukemia. 2010;24(7):1249–57. doi: 10.1038/leu.2010.104. PubMed PMID: 20520638.

113. Abate DA, Watanabe S, Mocarski ES. Major human cytomegalovirus structural protein pp65 (ppUL83) prevents interferon response factor 3 activation in the interferon response. J Virol. 2004;78(20):10995–1006. PubMed PMID: 15452220.

114. Browne EP, Wing B, Coleman D, Shenk T. Altered cellular mRNA levels in human cytomegalovirus-infected fibroblasts: viral block to the accumulation of antiviral mRNAs. J Virol. 2001;75(24):12319–30. PubMed PMID: 11711622.

115. Slobedman B, Stern JL, Cunningham AL, Abendroth A, Abate DA, Mocarski ES. Impact of human cytomegalovirus latent infection on myeloid progenitor cell gene expression. J Virol. 2004;78(8):4054–62. PubMed PMID: 15047822.

116. Chan G, Bivins-Smith ER, Smith MS, Smith PM, Yurochko AD. Transcriptome analysis reveals human cytomegalovirus reprograms monocyte differentiation toward an M1 macrophage. J Immunol. 2008;181(1):698–711. PubMed PMID: 18566437.

117. Mezger M, Bonin M, Kessler T, Gebhardt F, Einsele H, Loeffler J. Toll-like receptor 3 has no critical role during early immune response of human monocyte-derived dendritic cells after infection with the human cytomegalovirus strain TB40E. Viral Immunol. 2009;22(6):343–51. doi: 10.1089/vim.2009.0011. PubMed PMID: 19951172.

118. Zhu H, Cong JP, Shenk T. Use of differential display analysis to assess the effect of human cytomegalovirus infection on the accumulation of cellular RNAs: induction of interferon-responsive RNAs. Proc Natl Acad Sci U S A. 1997;94(25):13985–90. PubMed PMID: 9391139.

119. Boyle KA, Pietropaolo RL, Compton T. Engagement of the cellular receptor for glycoprotein B of human cytomegalovirus activates the interferon-responsive pathway. Mol Cell Biol. 1999;19(5):3607–13. PubMed PMID: 10207084; PubMed Central PMCID: PMCPMC84158.

120. Yurochko AD, Huang ES. Human cytomegalovirus binding to human monocytes induces immunoregulatory gene expression. J Immunol. 1999;162(8):4806–16. PubMed PMID: 10202024.

121. Preston CM, Harman AN, Nicholl MJ. Activation of interferon response factor-3 in human cells infected with herpes simplex virus type 1 or human cytomegalovirus. J Virol. 2001;75(19):8909–16. doi: 10.1128/JVI.75.19.8909-8916.2001. PubMed PMID: 11533154; PubMed Central PMCID: PMCPMC114459.

122. Netterwald JR, Jones TR, Britt WJ, Yang SJ, McCrone IP, Zhu H. Postattachment events associated with viral entry are necessary for induction of interferon-stimulated genes by human cytomegalovirus. J Virol. 2004;78(12):6688–91. doi: 10.1128/JVI.78.12.6688-6691.2004. PubMed PMID: 15163760; PubMed Central PMCID: PMCPMC416537.

123. Taylor RT, Bresnahan WA. Human cytomegalovirus immediate-early 2 protein IE86 blocks virus-induced chemokine expression. J Virol. 2006;80(2):920–8. doi: 10.1128/JVI.80.2.920-928.2006. PubMed PMID: 16378994; PubMed Central PMCID: PMCPMC1346867.

124. Fu YZ, Su S, Gao YQ, Wang PP, Huang ZF, Hu MM, et al. Human Cytomegalovirus Tegument Protein UL82 Inhibits STING-Mediated Signaling to Evade Antiviral Immunity. Cell Host Microbe. 2017;21(2):231–43. doi: 10.1016/j.chom.2017.01.001. PubMed PMID: 28132838.

125. Khan N, Bruton R, Taylor GS, Cobbold M, Jones TR, Rickinson AB, et al. Identification of cytomegalovirus-specific cytotoxic T lymphocytes in vitro is greatly enhanced by the use of recombinant virus lacking the US2 to US11 region or modified vaccinia virus Ankara expressing individual viral genes. J Virol. 2005;79(5):2869–79. PubMed PMID: 15709006.

126. Taylor RT, Bresnahan WA. Human cytomegalovirus IE86 attenuates virus- and tumor necrosis factor alpha-induced NFkappaB-dependent gene expression. J Virol. 2006;80(21):10763–71. PubMed PMID: 17041226.

127. Miller DM, Zhang Y, Rahill BM, Waldman WJ, Sedmak DD. Human cytomegalovirus inhibits IFN-alpha-stimulated antiviral and immunoregulatory responses by blocking multiple levels of IFN-alpha signal transduction. J Immunol. 1999;162(10):6107–13. PubMed PMID: 10229853.

128. Le VT, Trilling M, Wilborn M, Hengel H, Zimmermann A. Human cytomegalovirus interferes with signal transducer and activator of transcription (STAT) 2 protein stability and tyrosine phosphorylation. J Gen Virol. 2008;89(Pt 10):2416–26. doi: 10.1099/vir.0.2008/001669-0. PubMed PMID: 18796709.

129. Biolatti M, Dell’Oste V, Pautasso S, Gugliesi F, von Einem J, Krapp C, et al. Human Cytomegalovirus Tegument Protein pp65 (pUL83) Dampens Type I Interferon Production by Inactivating the DNA Sensor cGAS without Affecting STING. J Virol. 2018;92(6). doi: 10.1128/JVI.01774-17. PubMed PMID: 29263269; PubMed Central PMCID: PMCPMC5827387.

130. Marshall EE, Geballe AP. Multifaceted evasion of the interferon response by cytomegalovirus. J Interferon Cytokine Res. 2009;29(9):609–19. Epub 2009/08/28. doi: 10.1089/jir.2009.0064. PubMed PMID: 19708810; PubMed Central PMCID: PMC2743745.

131. Verma S, Benedict CA. Sources and signals regulating type I interferon production: lessons learned from cytomegalovirus. J Interferon Cytokine Res. 2011;31(2):211–8. doi: 10.1089/jir.2010.0118. PubMed PMID: 21226618; PubMed Central PMCID: PMCPMC3036178.

132. Goodwin CM, Ciesla JH, Munger J. Who’s Driving? Human Cytomegalovirus, Interferon, and NFkappaB Signaling. Viruses. 2018;10(9). doi: 10.3390/v10090447. PubMed PMID: 30134546.

133. van Boxel-Dezaire AH, Rani MR, Stark GR. Complex modulation of cell type-specific signaling in response to type I interferons. Immunity. 2006;25(3):361–72. doi: 10.1016/j.immuni.2006.08.014. PubMed PMID: 16979568.

134. Daffis S, Suthar MS, Szretter KJ, Gale M, Jr., Diamond MS. Induction of IFN-beta and the innate antiviral response in myeloid cells occurs through an IPS-1-dependent signal that does not require IRF-3 and IRF-7. PLoS Pathog. 2009;5(10):e1000607. doi: 10.1371/journal.ppat.1000607. PubMed PMID: 19798431; PubMed Central PMCID: PMCPMC2747008.

135. Browne EP, Shenk T. Human cytomegalovirus UL83-coded pp65 virion protein inhibits antiviral gene expression in infected cells. Proc Natl Acad Sci U S A. 2003;100(20):11439–44. PubMed PMID: 12972646.

136. Jahn G, Scholl BC, Traupe B, Fleckenstein B. The two major structural phosphoproteins (pp65 and pp150) of human cytomegalovirus and their antigenic properties. J Gen Virol. 1987;68(Pt 5):1327–37. doi: 10.1099/0022-1317-68-5-1327. PubMed PMID: 3033138.

137. Dobin A, Davis CA, Schlesinger F, Drenkow J, Zaleski C, Jha S, et al. STAR: ultrafast universal RNA-seq aligner. Bioinformatics. 2013;29(1):15–21. doi: 10.1093/bioinformatics/bts635. PubMed PMID: 23104886; PubMed Central PMCID: PMCPMC3530905.

